# Mineral-guided molecular enrichment: An interfacial driving force for protocell emergence on early Earth

**DOI:** 10.1101/2025.11.12.687952

**Authors:** Weifeng Chen, Rongfeng Zheng, Zhaosen Luo, Guoquan Wu, Jiayu Lei, Haochun Gou, Hao Li, Zhili Wang, Wenxuan Xiao, Chunhui Li, Guozhi Yu

## Abstract

The emergence of protocells from a dilute prebiotic environment is a fundamental challenge in origins-of-life research. which requires the simultaneous overcoming of molecular dilution, the establishment of metabolic cycles and the formation of selectively permeable compartments. But where and how such critical conditions are satisfied is very obscure and debated. We demonstrate, using contemporary model proteins and enzymatic systems, that mineral surfaces can facilitate the co-adsorption and spatial arrangement of nucleotides, proteins, and lipids into locally enriched, protocell-like assemblies under the examined laboratory conditions. We showed that geochemically relevant minerals efficiently co-adsorb and colocalize diverse biomolecules, creating crowded interfacial microenvironments that support enzyme cascades with substrate channeling-like behavior. Subsequently, these mineral-protein complexes template the assembly of lipid membranes, leading to the formation of discrete compartments that maintain metabolic activity while permitting molecular exchange. This mineral-guided surface enrichment (MSE) mechanism integrates key aspects of “metabolism-first” and “membrane-first” scenarios into a unified pathway, reconciling long-standing conceptual divides in prebiotic chemistry. Our findings establish mineral interfaces as active organizers of protocell emergence, offering a geochemically plausible framework for the origin of cellular life.

## Introduction

The transition from abiotic chemistry to Darwinian evolution in a cellular context remains one of the most fundamental and unresolved challenges in science. A critical prerequisite for origin of life is the emergency of molecularly crowded, compartmentalized systems capable of sustaining metabolic cycles and preserving informational polymers^1, 2, 3^. The primordial Earth, however, likely presented a vast and dilute aqueous environment, posing a fundamental thermodynamic paradox: the same medium that enables prebiotic chemistry also impedes the spontaneous assembly and interaction of macromolecules, this is also known as the dilution paradox ^4-6^. Although liquid-liquid phase separation (LLPS) offers one possible path to compartmentalization ^7-10^, it typically requires concentrations that are unlikely to have been widespread on the early Earth. Thus, a central challenge in origins-of-life research is to identify mechanisms that could overcome this thermodynamic hurdle—enabling the concentration of diverse molecules, supporting primitive metabolism, and enabling selective encapsulation—under plausible prebiotic dilution conditions ^11,12^.

This challenge is further reflected in the long-standing conceptual divide between “metabolism-first” ^13,14^ and “membrane-first” ^15-17^ scenarios. The former emphasizes the initiation of self-sustaining chemical networks, while the latter focuses on the early formation of boundary structures capable of maintaining chemical gradients. While recent hybrid models ^18^ have sought to integrate these views, a geochemically plausible framework that reconciles these perspectives under prebiotically relevant conditions has remained elusive. Mineral-water interfaces have long been recognized as crucial arenas for prebiotic chemistry, offering a potential solution to the dilution problem. It has long been recognized that mineral-water interfaces potentially offer a solution to the dilute paradox by selectively adsorbing and concentrating organic molecules, catalyzing the formation of biopolymers including peptides and nuclei acids^19-23^, generating electrochemical gradients which mediate the phase separation of polymers into distinct compartments^24^, and introducing non-equilibrium processes ^25^ through wet-dry cycles at these interfaces. However, such individual functions have never been integrated in a unified framework where the molecular enrichment, proto-metabolism^26, 27^, and compartmentalization can be realized in a single and continuous pathway ^28, 29^.

Here we show that mineral interfaces act as dynamic platforms that orchestrate the hierarchical assembly of protocellular structures, directly driving their emergence. We propose a mechanism of mineral-guided surface molecular enrichment (MSE), where geochemically relevant mineral surfaces first mediate the co-adsorption and spatial organization of key biomacromolecules. This process creates a crowded two-dimensional reaction space, establishing the conditions necessary for the emergence of surface-bound, proto-metabolic pathways. Subsequently, these functionalized protein-mineral complexes serve as templates for the assembly of lipid membranes, leading to the formation of discrete, catalytically active compartments.

Our findings reveal that mineral surfaces not only concentrate diverse biomolecules but also support the spatial organization of multi-enzyme cascades, creating a configuration that facilitates coupled catalytic reactions in a process analogous to substrate channeling in biological cells. This synergistic pathway, which integrates molecular crowding, interfacial catalysis, and membrane formation, delineates a robust and plausible route from disorganized molecules in a prebiotic soup to functionally integrated, protocellular systems. Our work thus establishes mineral surfaces as foundational and active platforms for prebiotic systems chemistry, offering a coherent mechanism that couples interfacial metabolic activity with hierarchical self-assembly under conditions relevant to the early Earth.

## Results

### Mineral-mediated stable co-enrichment of different types of biomolecules

To probe how geochemical interfaces could have organized prebiotic matter, we examined the ability of mineral particles to concentrate and co-localize DNA, RNA and protein. Considering reports indicating that relatively unstructured protein segments are preferentially adsorbed by silica ^30-32^, and that primordial amino-acid condensation likely produced intrinsically disordered sequences ^33^, we performed all experiments using two fusion constructs that meet these criteria: FUS-IDR-sfGFP^34^ and BFP-hnRNPA1-IDR^35^ (sequences detailed in Supplementary Table 1). Silica microspheres adsorbed Cy3–DNA, FITC–RNA and BFP–tagged protein, each detected in its respective fluorescence channel (Fig. 1a–c), and—crucially—supported the simultaneous co-adsorption of all three biomolecular classes at their surfaces. Quantitative fluorescence mapping and line-scan analysis revealed tight spatial overlap of the three signals at the silica interface (Fig. 1d–e; Supplementary Fig. 3), consistent with the formation of locally enriched microenvironments that could concentrate reaction substrates and intermediates to promote surface-confined reaction fluxes akin to protometabolic cycles. Furthermore, the adsorbed material was also proven resilient to environmental flux: after repeated washing (10 times), individual FUS-IDR-GFP molecules and ternary protein – RNA – DNA assemblies remained detectable on CaCO_3_, montmorillonite and SiO_2_ (Fig. 1f, g). Crucially, this co-adsorption is highly stable against physical perturbation (Fig. 1f, g), demonstrating that these enriched microenvironments could persist within the dynamic conditions of the early Earth. Together, these observations identify diverse mineral surfaces as robust scaffolds for resisting dilution and for creating spatially distinct, semi-stable niches that could scaffold early, surface-mediated proto-metabolic and compartmentalization processes. Extending this, we further tested that elevated temperature (70 ° C)-plausible condition on early earth-further enhanced protein adsorption across all tested minerals (Supplementary Fig. 1), with montmorillonite showing the highest capacity owing to its layered, negatively charged architecture, suggesting that minerals could have simultaneously concentrated and stabilized primordial biomolecules under geologically relevant conditions on early Earth.

**Figure 1.**
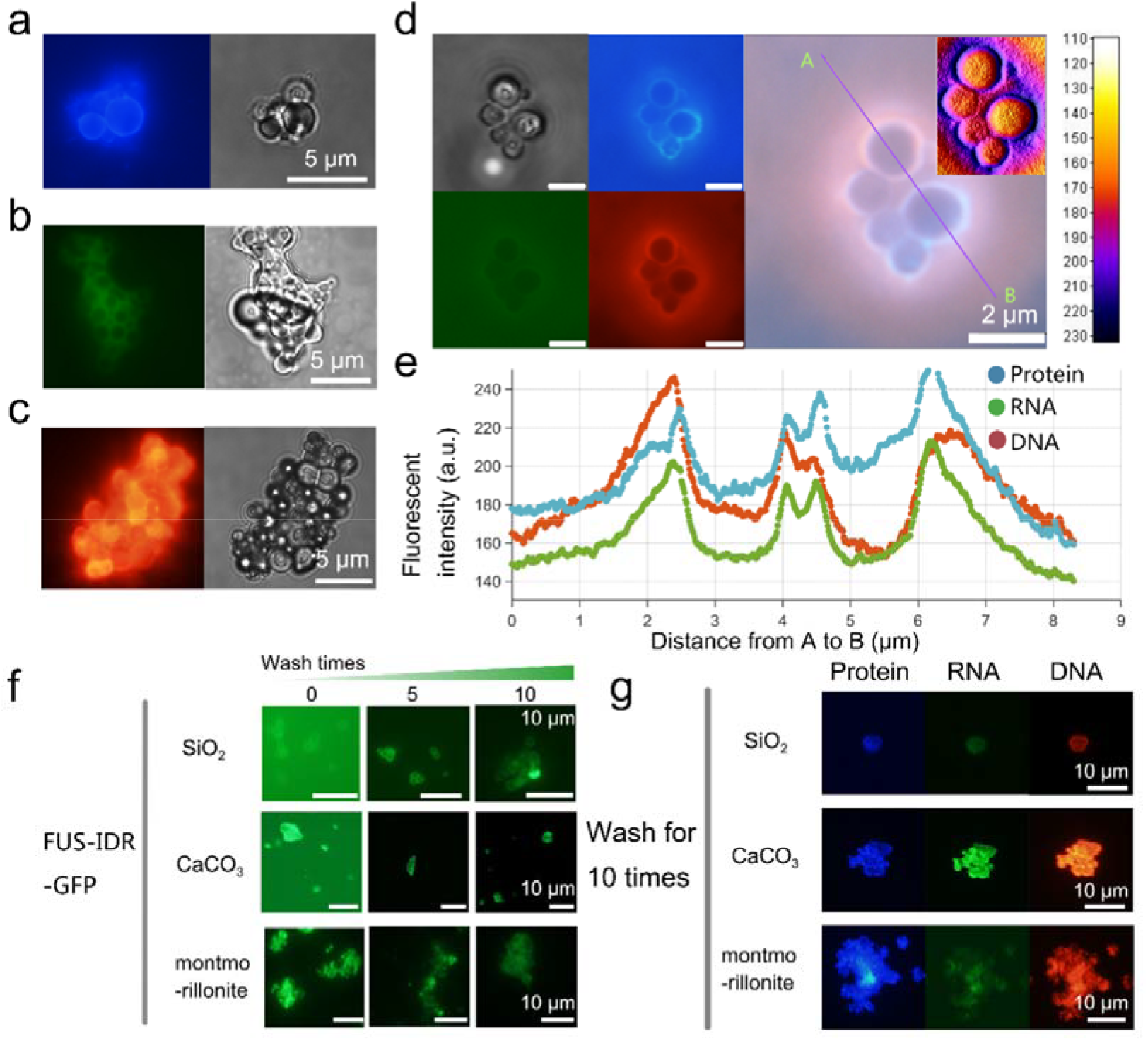
Co-adsorption, spatial co-localization, and retention stability of biomacromolecules mediated by minerals. a-c, Single-channel fluorescence images of individually labeled biomolecules adsorbed on silica particles: (a) BFP-hnRNPA1-IDR protein (blue), (b) FITC-labeled RNA (green), and (c) Cy3-labeled DNA (red). The right panel shows corresponding bright-field images. d, Composite analysis images of co-adsorbed biomolecules: from left to right, bright-field, individual fluorescence channels (protein in blue; RNA in green; DNA in red), merged fluorescence image, and an RGB integrated fluorescence heatmap inset (color bar represents cumulative intensity, a.u.). e, Fluorescence intensity profiles along the path A-B in d (protein in blue; RNA in green; DNA in red), showing co-localization of the three biomacromolecules. f, Retention of FUS-IDR-GFP fluorescence (green) on SiO_2_, montmorillonite, and CaCO_3_ mineral surfaces washed 0, 5, and 10 times. Each row of images was taken with the same acquisition parameters. g, Co-adsorption and retention of proteins (blue), RNA (green), and DNA (red) on specified mineral surfaces after 10 times washing. Scale bars are showed in each subfigures.

### Heterogeneous molecular layering and limited fluidity on mineral surfaces

After analyzing the adsorption of three types of biomacromolecules onto different mineral particles, we investigated the adsorption of proteins onto silica particles as a case study to further delineate the physical characteristics of MSE. We started with a quantitative investigation of the protein shell distribution on the surfaces of spherical silica particles. Figure 2.a depicts the coordinate system alongside additional parameters (see to Supplementary Note 1 for mathematical details). Figure 2b illustrated that the radial fluorescence intensity profile was deconvoluted with a dual-exponential decay model to quantify the spatially structured molecular layers, expressed as *I(r) = α*. *exp(-λ*_*1*_*r) + β*. *exp(-λ*_*2*_*r) + bg*. This model delineates two distinct physical regimes: (i) a near-field component characterized by amplitude α and decay rate λ_1_, representing a dense layer of FUS-IDR-GFP firmly adhered to the mineral surface; and (ii) a far-field component defined by amplitude β and decay rate λ_2_, corresponding to a more diffuse corona of weakly interacting molecules in equilibrium with the bulk solution. We examined the parameter space across different macromolecular concentrations to comprehend the distribution of various parameters (Fig. 2c). The radar plot results demonstrate that, throughout both high- and low-concentration regimes, α, β, λ_1_, and λ_2_ exhibit extensive variability, but the background (bg) stays distinctly separate (the average bg for high and low concentration groups are 55.04 and 21.08, a.u., respectively). This seemingly unreasonable parameter distribution indicates a credible scenario: significant particle heterogeneity throughout the MSE system. This variety is likely resulted from differences in inorganic particle geometry and surface characteristics, random fluctuations in local bulk concentrations, and interactions among adjacent particles, therefore providing biomacromolecules with a wide range of ecological niches.

**Figure 2.**
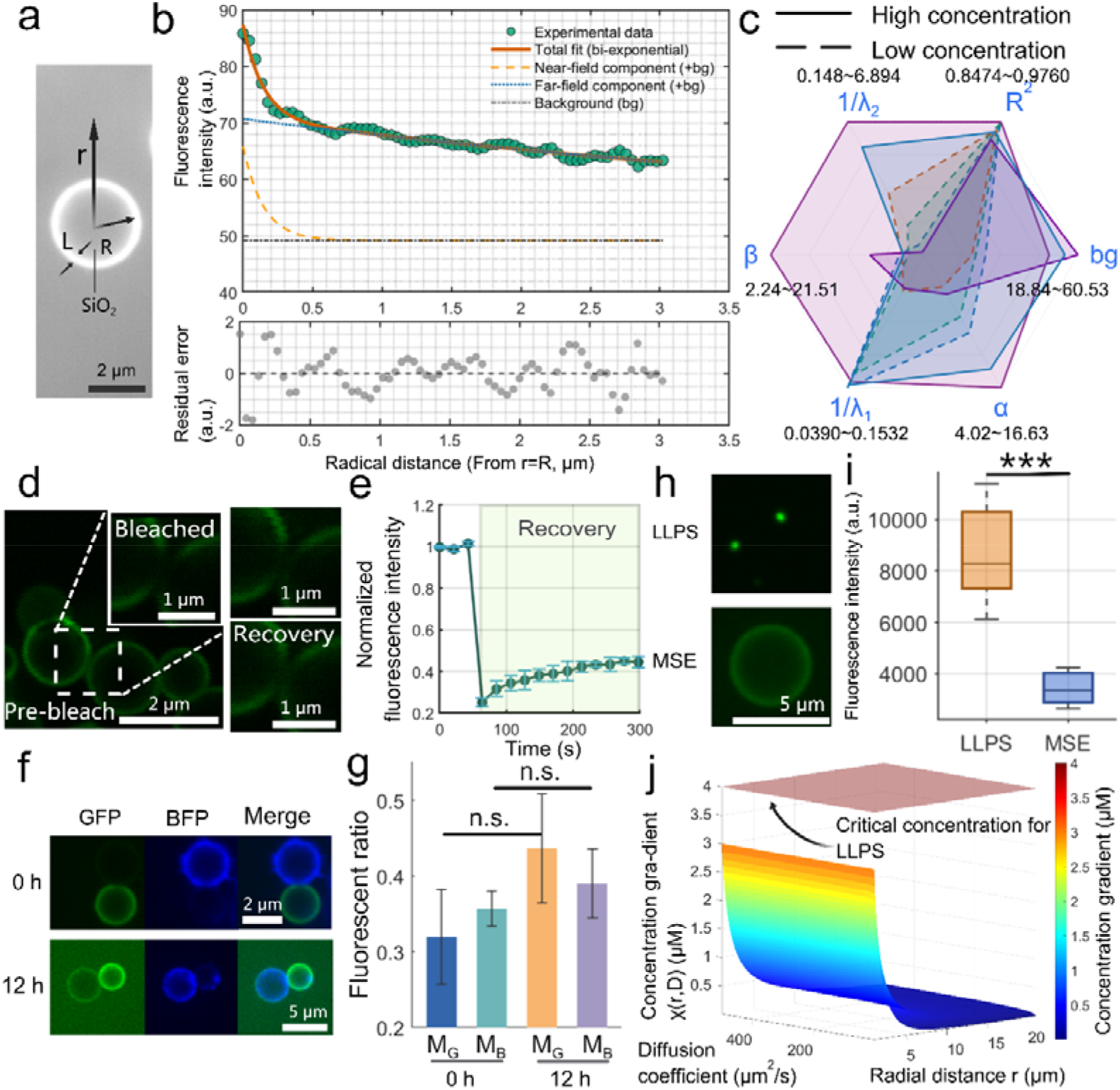
Characterization of the physical properties of molecular enrichment on mineral surfaces. a, Schematic representation of the radial coordinate system for a spherical silica particle with radius *R*. The shell thickness (*L*) and distance from the surface (*r*) define the spatial parameters of protein dispersion. b, The radial fluorescence intensity distribution was fitted using a double exponential decay model:. The top panel shows experimental data points and the model fit curve (R^2^ = 0.9760), and the bottom panel shows a residual plot indicating a random error distribution. c, Radar plot of model fitting parameters at different protein concentrations. Three representative samples were chosen for each concentration group. d-e, Time-lapse images (d) and quantitative analysis (e) of fluorescence recovery after photobleaching (FRAP) of a FUS-IDR-GFP protein corona on a silica particle surface, showing an approximately 20% recovery in fluorescence intensity over 300 seconds. Error bars represent the standard deviation of three repeated experiments. f, Fusion kinetics of pre-washed FUS-IDR-GFP / BFP-hnRNPA1-IDR labeled protein-silica particles over 12 hours. g, Mander’s coefficient (M_B_/M_G_) quantitatively analyzes the degree of molecular intermixing during fusion. Data are presented as mean ± s.d. (n = 3 pairs of particles per group, unpaired two-tailed Student’s *t*-tests, the p-values for the M_G_ and M_B_ groups were 0.10 and 0.32, respectively). h-i, Fluorescence images (h) and quantitative analysis of fluorescence intensity (i) of liquid-liquid phase separation (LLPS) condensates (top) and MSE (bottom) under identical conditions, confirming that LLPS has a higher molecular enrichment efficiency (\*\**p* < 0.001; unpaired two-tailed *t*-test; n = 20 per group). j, 3D simulation of a concentration gradient formed by diffusion (along the radial axis r). Scale bars are showed in each subfigures.

To elucidate the origin of surface permeability of the interfacial protein corona, we employed fluorescence recovery after photobleaching (FRAP) to evaluate the fluidity of the FUS-IDR-GFP corona. Post-photobleaching, the fluorescence of surface shells was seen to recover 20% of its initial intensity over 300 seconds—a slight yet noteworthy fluidity (Fig. 2d-e). To address potential hydrodynamic flushing in early-Earth environments, we introduced buffer rinsed GFP[SiO_2_] and BFP[SiO_2_] and measured their fusion rates. Contact fusion studies with GFP/BFP-coated silica particles demonstrated prolonged intermixing between neighboring compartments for 12 hours (Fig. 2f). While the overall degree of molecular intermixing, as measured by Mander’s coefficients (M_G_, M_B_), did not show a statistically significant change over 12 hours (p > 0.05, Fig. 2g), we observed a slow redistribution of fluorescent proteins between adjacent particles. This suggests that the protein corona, while largely stable, possesses a limited fluidity. This combination of long-term stability and slow short-range dynamics is functionally critical: it allows for the preservation of distinct molecular identities in neighboring compartments, a prerequisite for functional specialization, while still permitting the interfacial molecular movement necessary for proto-metabolic processes. This dynamic activity validates the mineral interface’s ability to facilitate substrate movement crucial for protometabolic continuity. Simultaneously, the slow yet persistent macromolecular migration preserves heterogeneity within the inorganic-particle surface layer, establishing a foundation for functionally separate metabolic compartments analogous to modern subcellular structures—or even tissues.

Prompted by fluidity and fusion dynamics, we investigated whether MSE could complement liquid–liquid phase separation (LLPS) in dilute prebiotic environments. Fluorescence intensity comparisons revealed LLPS condensates exhibited significantly higher biomolecular sequestration than MSE (Fig. 2h, i). MSE’s comparatively diffuse enrichment profile (Fig. 2i) confirms its intrinsically weaker sequestration efficiency—a direct consequence of its reliance on surface adsorption rather than bulk demixing. Paradoxically, this apparent “limitation” resolves a prebiotic paradox: in primordial settings where biomolecule concentrations were too low for spontaneous LLPS nucleation ^36^, MSE provided a critical pre-concentrating step.

To rationalize the interfacial enrichment observed experimentally, we computed the steady-state radial concentration field around a spherical mineral particle under linearized adsorption–desorption kinetics (Supplementary note. 1). For the general spherical solution of the linear reaction–diffusion equation the concentration decays as 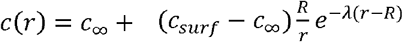, where *R* is the particle radius, *c*_∞_ the bulk concentration, and 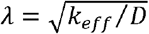 is the inverse penetration length with *k*_eff_ *= k*_off_ *+ k*_on_ *c*_∞_ (the adsorption coefficient *k*_on_ is expressed in µM^-1^·s^-1^ and *k*_off_ in s^-1^; diffusion coefficients *D* are in µm^2^·s^-1^). The *R/r* prefactor enforces the geometric dilution in spherical coordinates and recovers the planar exponential form when *r ≈ R* and *L « R*. The computed map in Fig. 2j uses this spherical solution to show *χ(r, D) = c(r) - c*_∞_ across diffusion coefficients and radial distances (representative values *c*_*surf*_ *=4 µM, c*_∞_ *= 1 µM, c*_*crit*_ *= 5 µM* were used for illustration). For these parameters the surface remains below the bulk LLPS threshold across the explored domain; however, modest changes in boundary values (for example increases in *c*_*surf*_, reductions in *c*_*crit*_, or changes in adsorption/desorption kinetics, mimicking the fluctuations of plausible cases) can move the near-surface region above the critical manifold and permit interfacial nucleation (Supplementary Fig. 3). Note that increasing *D* increases the penetration length *1/λ*, thereby extending the radial range where *c(r)* may approach the threshold, but does not by itself increase *c*_*surf*_ — therefore the combined effect on LLPS nucleation depends jointly on the boundary concentration and the penetration length.ackn

### Silica-scaffold-supported metabolic activity

Building upon the observed fluidity of the interfacial protein corona, we next investigated whether these dynamic mineral scaffolds could organize and support functional, multi-step biochemical reactions, a prerequisite for proto-metabolism. To probe this, we assembled a two-step enzyme cascade on silica microspheres. The system comprised glucose oxidase (GOx) and horseradish peroxidase (HRP), which act sequentially to convert glucose into H_2_O_2_ and then utilize the H_2_O_2_ to oxidize Amplex Red into the fluorescent product, resorufin ^37 38^. This spatial juxtaposition is known to enhance overall reaction efficiency by minimizing the diffusion of intermediates, a process analogous to substrate channeling in contemporary cells ^39^. We co-localized GOx and HRP onto the silica surfaces along with two distinct IDR-fused fluorescent proteins (FUS-IDR-GFP and hnRNPA1-IDR-BFP), which served to delineate the reactive surface and promote enzyme retention. After removing unbound enzymes via washing, fluorescence microscopy confirmed the stable co-localization of the fluorescent marker proteins on the particle surfaces. Crucially, the robust emission from resorufin, localized exclusively at the mineral interface, provided direct evidence of locally coupled catalytic activity. These findings establish that mineral interfaces can serve as effective scaffolds for assembling synergistic, proto-metabolic networks, thereby creating a foundation for the subsequent evolution of compartmentalized protocellular systems. To quantitatively assess the kinetic effects of interfacial confinement, we conducted stochastic simulations of the two-enzyme cascade under both bulk and mineral-guided surface enrichment (MSE) conditions (Fig. 3c; Supplementary Note 2). The findings demonstrated a concentration-dependent transition in catalytic efficacy. In high enzyme concentrations, bulk systems surpassed MSE because of diffusion limitations and fast substrate exhaustion at the surface. At moderate concentrations, MSE demonstrated a significant initial acceleration phase—indicative of effective substrate capture— subsequently transitioning to a gradual product accumulation when the localized substrate reservoir was depleted. Notably, at low enzyme concentrations, MSE consistently exhibited superior reaction speeds and total yields compared to bulk, indicating that surface enrichment confers a kinetic advantage specifically in the dilute regimes pertinent to prebiotic circumstances. These findings demonstrate that MSE not only immobilizes enzymes but also actively alters the reaction landscape, facilitating localized metabolic continuity despite limited global reactant availability.

**Figure 3.**
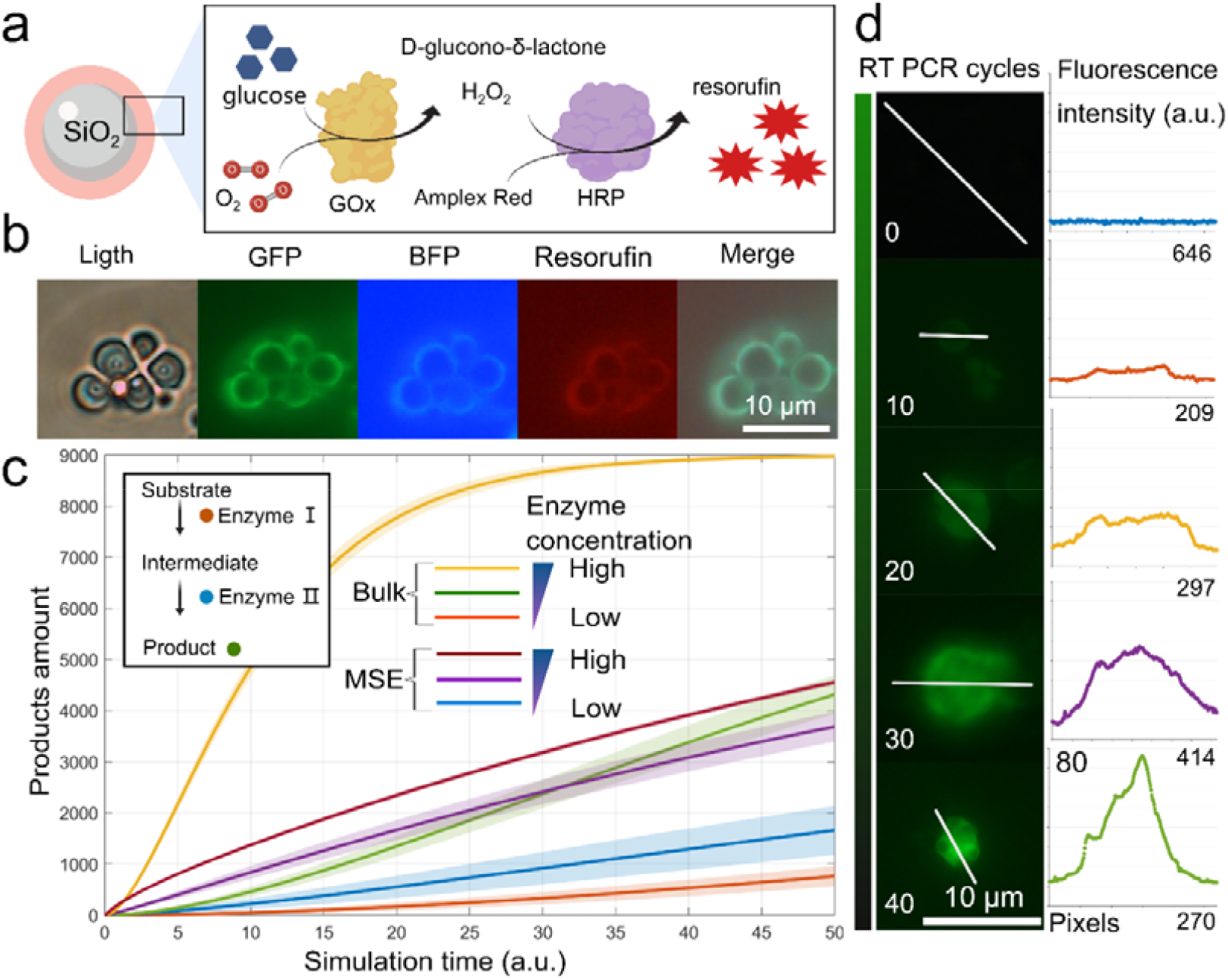
Enzyme cascade assembly and imaging on silica particles. a, Schematic diagram of the glucose oxidase (GOx) and horseradish peroxidase (HRP) cascade reaction. GOx catalyzes the conversion of glucose to D-glucono-δ-lactone and hydrogen peroxide, and HRP subsequently uses the hydrogen peroxide to oxidize Amplex Red into the fluorescent product resorufin (red star). b, Microscopic images (from left to right): bright-field (light), FUS-IDR-GFP, BFP-hnRNPA1-IDR, resorufin fluorescence (red), and merged channel images. The images show that the FUS-IDR-GFP / BFP-hnRNPA1-IDR signal and resorufin synthesis co-localize on the silica surface, confirming the immobilization and functional coupling of the proteins. c, Stochastic simulations comparing bulk and MSE environments across enzyme concentration regimes. At high enzyme concentrations, bulk reactions yield higher overall product levels; at intermediate concentrations, MSE displays an initially faster but later plateauing output; and at low concentrations, MSE sustains greater catalytic throughput, highlighting the confinement advantage under dilute conditions. Shaded regions denote standard deviations across replicate simulations (n = 200 for each group). d, Reverse transcription PCR in the presence of silica particles. The left image sequence shows the increasing intensity of SYBR Green I fluorescence with the number of cycles, all images acquired with the same parameters. The right graph shows a histogram of light intensity scans along the equal-length straight lines in the top images (The horizontal axis denotes the number of pixels and, the vertical axis denotes the fluorescence intensity, a.u.). Scale bars are showed in each subfigures.

Given that the primordial soup likely contained both nucleic acids and proteins, we further sought to determine if such mineral interfaces could also support enzymatic processes involving nucleic acids. We performed a reverse transcription-polymerase chain reaction (RT-PCR) assay directly within a suspension of silica particles, monitoring the amplification of a target RNA template via SYBR Green I fluorescence. Time-lapse microscopy and quantitative fluorescence analysis revealed a progressive, cycle-dependent intensification of the SYBR Green I signal localized on the silica surfaces. This demonstrates that the enzymatic machinery for both reverse transcription and DNA amplification remains functional when adsorbed onto the mineral particles, suggesting these surfaces could have supported not only primordial metabolism but also rudimentary forms of macromolecule replication and transfer.

### Mineral-templated formation of metabolically active protocell-like compartments

After establishing that mineral surfaces facilitate enzyme colocalization and functional coupling (Fig. 3), we investigated whether these pre-organized molecular architectures could template higher-order structural complexity — specifically, the formation of spatially confined lipid–protein assemblies capable of supporting metabolic compartmentalization. In control systems comprising bare silica particles and palmitoleic acid (PAL), lipid assembly primarily resulted in free lipid droplets (Fig. 4b, arrowheads), with occasional adsorption onto silica surfaces (arrows), indicating baseline interfacial activity. Introduction of FUS-IDR-GFP protein coronas markedly altered lipid distribution: pre-coated silica particles templated hybrid assemblies wherein lipids co-localized with protein coronas (Fig. 4c, arrows), though residual free droplets persisted (arrowheads), reflecting a dynamic equilibrium between surface-bound and mobile lipid phases.

**Figure 4.**
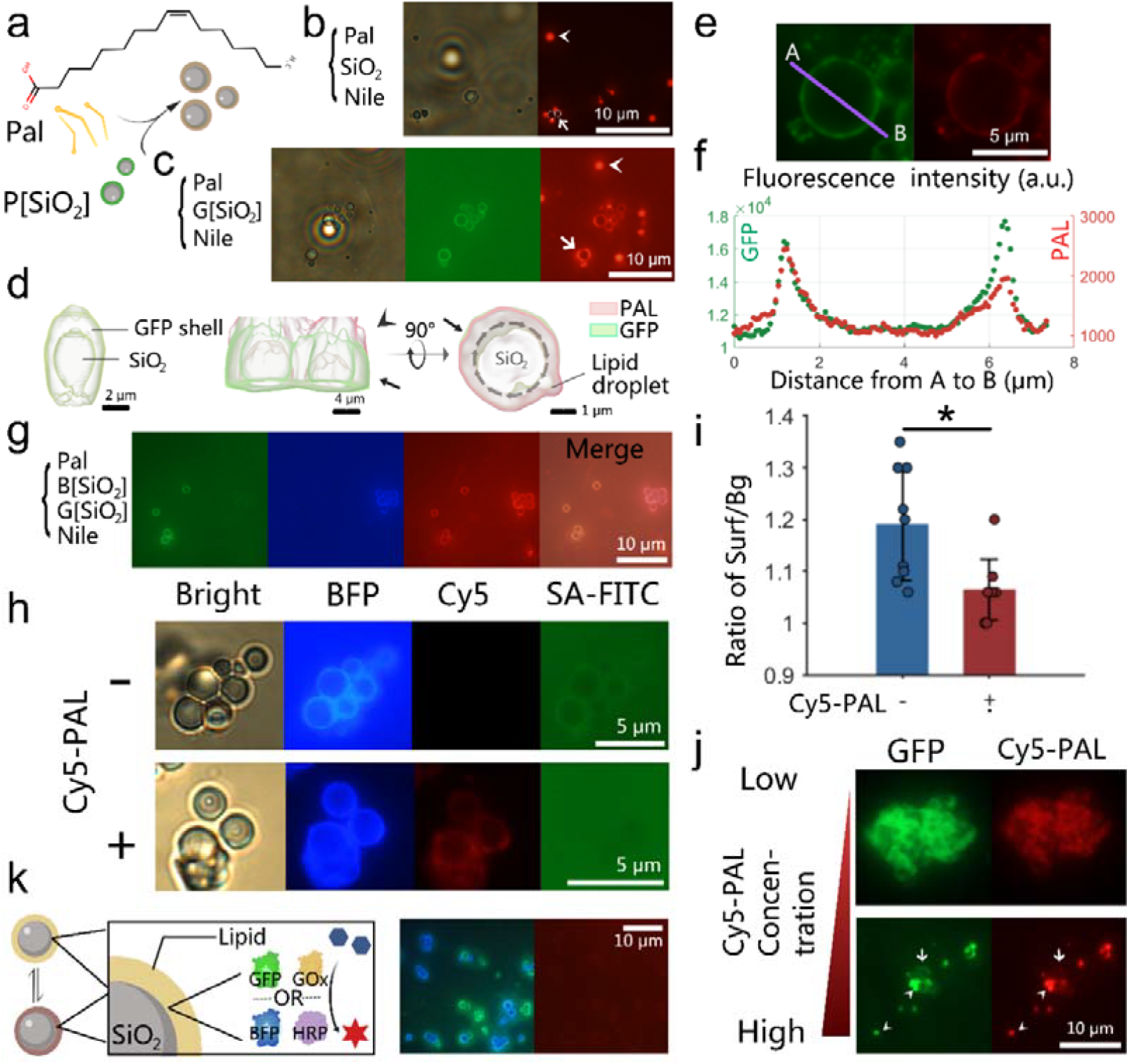
Hierarchical assembly of protocell-like compartments through silica-protein-lipid synergy. a, Schematic diagram of the experimental process related to b and c, where the chemical formula represents the palmitoleic acid (PAL) used in the experiment. b, Control system (silica + PAL). The left image is a bright-field image of a silica particle with a small amount of adsorbed lipid droplets. The right image is the corresponding Nile Red fluorescence (red) image, highlighting both free lipid droplets (arrow) and adsorbed lipid droplets (arrowhead). c, Protein-guided lipid assembly (silica + GFP + PAL). From left to right: bright-field image; GFP fluorescence (green) confirming the formation of a uniform protein corona; Nile Red channel (red) showing co-localization of the lipid membrane with the protein-coated particle (arrowhead), with some free droplets still present (arrow). d, 3D reconstructed image from optical sections, showing lipid-protein co-localization (arrowhead) and free lipid droplets (arrow). Left panel, GFP + silica; middle and right panel, GFP + PAL + silica. e-f, Co-localization analysis of GFP and PAL on the same silica particle surface. f is the quantitative intensity profile along segment AB in e. g, Compartmentalized hybrid system with mixed GFP+BFP assemblies. From left to right: GFP (green) and BFP (blue) labeled compartments remain separate; merged image confirms that a continuous lipid envelope encapsulates distinct protein-mineral cores. h, Permeability assay using FITC-streptavidin on silica-BFP particles with (+) or without (-) attached Cy5-PAL. i, Quantitative analysis of the results in h (n = 9 per group, independent two-sample t-test, *p* < 0.05, mean±s.d.). j, Interaction of GFP and Cy5-PAL with silica particles at different molar ratios. At low ratios, the two components spread on the silica surface to form a quasi-uniform shell (arrowhead); at high ratios, they co-localize outside the particle to form a dense, solid-state condensate (arrow). k, Metabolic coupling on lipid-encapsulated SiO_2_ particles. The left diagram is a schematic of the enzyme-coupled hybrid assembly, and the right image shows that resorufin fluorescence (red) synthesis is selectively localized on the SiO_2_ surface with both enzymes. Scale bars are showed in each subfigures.

**Figure 5.**
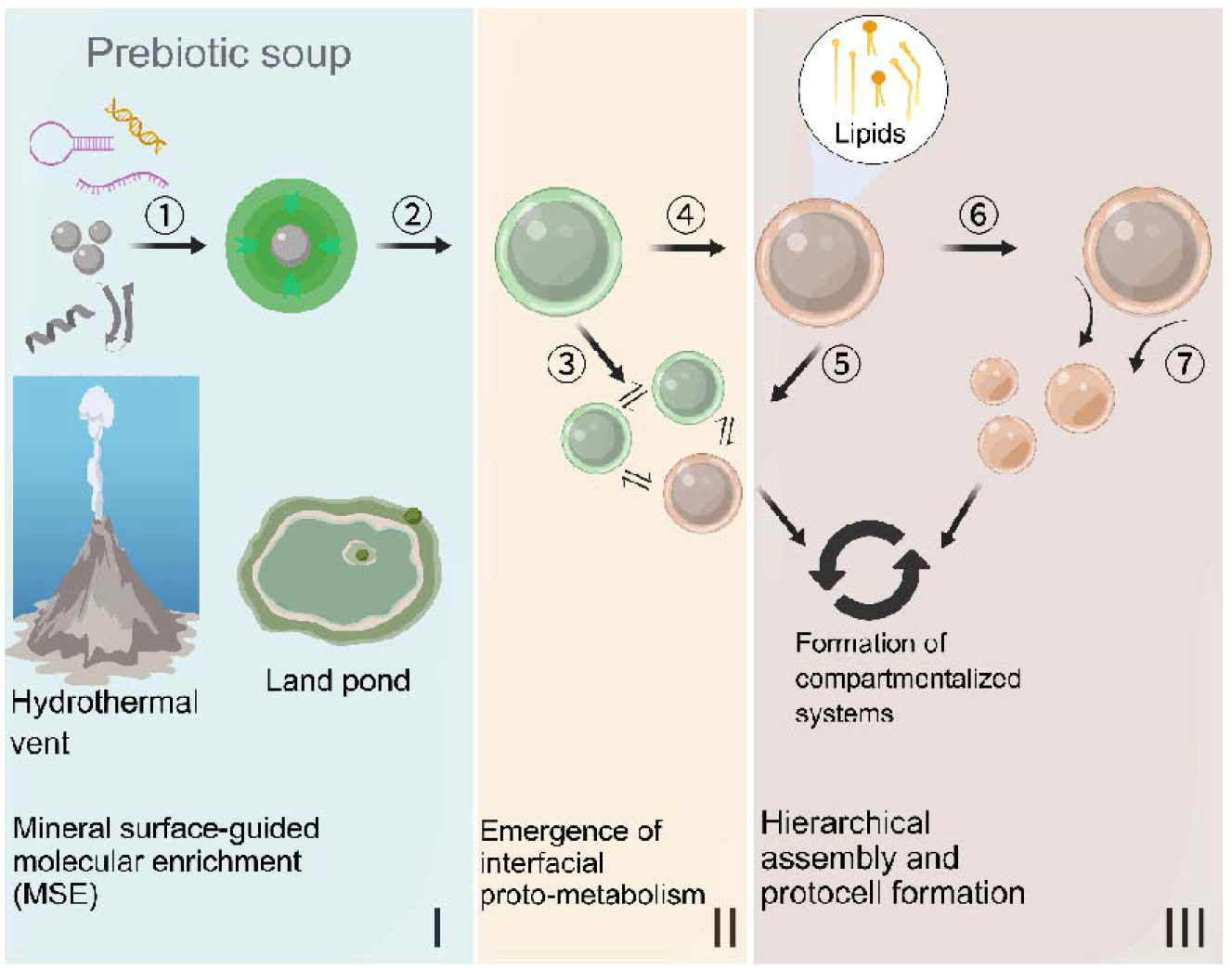
An integrated model for mineral-guided surface enrichment support for protocell emergence. (I) Mineral-guided surface molecular enrichment (MSE). In prebiotic environments such as hydrothermal vents or land ponds, essential biomolecules including nucleic acids and peptides co-adsorb onto mineral surfaces (1), concentrating them and overcoming the dilution problem inherent to the prebiotic soup. (II) Emergence of interfacial proto-metabolism. The resulting molecularly crowded interface facilitates enzyme cascades (2), establishing a proto-metabolic system analogous to substrate channeling. This system maintains dynamic exchange with other functionalized particles (3) or with ensuing lipid-coated assemblies (4). (III) Hierarchical assembly and protocell formation. Lipids subsequently assemble around the protein-mineral composites, templating the formation of a membrane boundary (5). This process enables the selective encapsulation of distinct catalytic cores (6), giving rise to compartmentalized systems that maintain functional heterogeneity and metabolic continuity, ultimately leading to the formation of a protocell (7).

Three-dimensional reconstructions revealed structural details of these complexes, illustrating clear lipid – protein co-localization (arrow) alongside persistent free lipid droplets (arrowhead, Fig. 4d). Quantitative colocalization analysis confirmed significant overlap between GFP and PAL signals (Fig. 4e–f). We further assessed the functional implications of these hybrid assemblies on molecular permeability: silica particles functionalized with Cy5-PAL exhibited reduced uptake of FITC–streptavidin compared to lipid-free controls (Fig. 4h– i), with an approximately 12.5% decrease in normalized fluorescence intensity (p < 0.05), consistent with modest barrier formation.

To probe compartmentalization dynamics, we amalgamated GFP- and BFP-coated assemblies for 2 hours (Fig. 4g). Lipid membranes selectively encapsulated distinct protein–mineral complexes, yielding compartments that maintained separate GFP or BFP identities—evidence that lipid layers serve as barriers minimizing surface-protein mixing. This templating effect underscores the role of mineral–protein cores in guiding lipid assembly toward spatially segregated, non-fusing protocell-like entities.

Upon lipid coating, the mineral-associated enzyme assemblies retained measurable catalytic activity, although the overall turnover rate decreased compared with uncoated samples (Fig. 3b). While this attenuation has been attributed to restricted substrate diffusion through the lipid layer (Fig. 4h, i), alternative factors may also contribute. In particular, direct lipid–enzyme interactions or subtle conformational changes could modulate enzyme accessibility and dynamics at the interface. Regardless of the underlying mechanism, the persistence of detectable catalysis within these semi-permeable shells indicates that the encapsulated systems remain functionally competent, providing a minimal model for protocell-like compartments capable of sustaining surface-confined reactivity under bounded conditions.

We additionally investigated the interaction between protein coronas and various phospholipids. Supplementary Fig. 5a illustrates the co-localization of phosphatidylserine (PS) and phosphatidylcholine (PC) with BFP-hnRNPA1-IDR on silica, highlighting the universality of lipid–protein interactions at mineral surfaces. Furthermore, as depicted in Supplementary Figure 5b, suspensions comprising PAL, silica, and Nile Red exhibited distinct stratification in the absence of protein (left tube). In contrast, the inclusion of BFP or GFP resulted in stable, homogeneous suspensions (middle and right tubes), thereby indicating that protein coronas facilitate lipid–mineral integration and stabilize heterogeneous mixtures—essential characteristics for prebiotic compartmentalization.

## Discussion

Our results delineate a mineral-guided strategy for protocell emergence that directly tackles the persistent issues of molecular dilution and uncontrolled compartmentalization. Prior studies emphasized the efficiency of bulk liquid–liquid phase separation (LLPS) as a mechanism for biomolecular condensation ^40,41^. However, such processes typically require micromolar to millimolar concentrations ^42-45^ seldom attainable in prebiotic settings. By contrast, we show that mineral-guided surface enrichment (MSE) generates interfacial microenvironments that locally lower the threshold for phase transitions. This divergence from bulk LLPS highlights how dimensional reduction at mineral interfaces creates a distinct physicochemical regime—less efficient in sequestration, yet particularly adapted to dilute primordial conditions.

Mineral-mediated organization has long been implicated in origins-of-life chemistry. Clay and silica surfaces are known to adsorb and even catalyze the polymerization of nucleotides and peptides ^23, 46^, while recent biophysical studies demonstrate that charged or wet-dry cycled interfaces can promote molecular crowding and phase transitions ^47^. Our observation that MSE co-localizes nucleic acids, proteins, and lipids under dilute conditions aligns with prior observations and offers experimental evidence that mineral surfaces function as dynamic controllers of molecular assembly rather than merely passive scaffolds. Furthermore, silica-bound protein coronas reduce the entropic cost of lipid ordering, consistent with models suggesting that pre-existing scaffolds promote amphiphile self-assembly ^48^, analogous to cytoskeletal templating in modern cells.

Our finding combine conflicting protocell scenarios into a continuous, mineral-directed pathway. Hybrid protocell systems have demonstrated that coacervates can be cloaked by fatty acid layers, yielding semi-permeable vesicles with selective uptake. We show that a similar transition can occur at mineral interfaces, where lipid encapsulation of protein–mineral composites yields distinct, metabolically active compartments. This process connects surface-anchored complexes to autonomous vesicles, so reconciling the seeming opposition between the “membrane-first” ^15-17^ and “metabolism-first” ^13,14^ models. Moreover, by sustaining enzyme cascades reminiscent of substrate channeling and supporting nucleic acid replication on surfaces, MSE offers a framework that couples enrichment, metabolism, and compartmentalization within geochemically plausible contexts.

Taken together, our work demonstrates that mineral interfaces are active organizers of protocell emergence—contrasting with bulk-phase models, supporting prior observations of interfacial chemistry, and extending the field by unifying disparate scenarios into a coherent pathway. This mineral-mediated hierarchy provides a versatile platform for systems chemistry on the early Earth and suggests testable routes for reconstructing the origins of cellular complexity. Additionally, it is essential to understand that our experiments use contemporary biomolecules and commercial buffers to probe physical mechanisms (adsorption, templating, lipid assembly). While the observed phenomena demonstrate plausible routes by which mineral surfaces could organize matter, further work with prebiotically plausible peptides, simple amphiphiles and primitive salts is required to assess whether identical behaviours would emerge under realistic early-Earth chemical conditions.

## Methods

Extraction and Preparation of Fluorescent Proteins The plasmids of the two fluorescent proteins (for the sake of simplicity, all GFP in this article represents FUS-IDR-GFP, and all BFP represents EBFP-hnRNPA1-IDR) were separately transformed into the E. coli BL21(DE3) strain. The transformed bacteria were cultured in LB medium containing kanamycin at 37 °C. When the optical density (OD) value reached 0.4-0.6, isopropyl β-D-1-thiogalactopyranoside (IPTG) (with a final concentration of 0.3 mM, Beyotime Biotechnology) was added, and the bacteria were then incubated with shaking at 28 °C for 8h. Recombinant bacterial cells expressing fused-proteins were pelleted by centrifugation (5,000×g, 10 min, 18°C). Pellets were resuspended in non - denaturing lysis buffer (pH=8.0) and disrupted via sonication (10 s on/off cycles, 300W, 20min). Clarified lysates(centrifuged for 30min at 4°C,12000g) were loaded onto nickel-chelated agarose columns (Beyotime Biotechnology) pre-equilibrated with non - denaturing lysis buffer (pH=8.0), followed by 0.5 h incubation at 4°C for protein binding. The column underwent sequential washing (5-column volumes, non - denaturing washing buffer (pH=8.0) and elution (non - denaturing elution buffer (pH=8.0). To recover target proteins, purified proteins were dialyzed overnight at 4°C against storage buffer to exchange buffer components. The concentration of proteins was determined using a BCA kit (Beyotime Biotechnology). See protein sequences for table S1. And the formulations of various buffer solutions are shown in supplementary table 4.

### Adsorption of Different Bio-Macromolecules on the Surfaces of Various Minerals

In both Figure 1 and Figure S1, all minerals, DNA, and RNA were dissolved in a 50 mM tris-HCL (pH 7.4) buffer solution, while proteins were dissolved in a buffer solution containing 50 mM Tris-HCl and 125 mM NaCl (pH 7.4). In the corresponding experiments, the final concentration of proteins (the sequence information is shown in Supplementary Table 1) was 0.5 mg/ml. The final concentration of DNA (18S rRNA Cy3 probe, Beyotime Biotechnology) and RNA (random sequence, 5’-FITC labeled CAACGACUCGCGCUUAAUUA, synthesized by Sangon Biotech) was 25 μg/ml. The final concentration of various minerals was 2.5 mg/ml. Unless otherwise specified, all microscopic photographs were taken using an Olympus BX53 microscope.

### Contact silica particles coated with different protein shells and analyze protein migration

Prepare separate stock solutions at final concentrations of 0.5 mg/mL protein (GFP or BFP) and 5 mg/mL silica suspension. After 5 min of vortex-mixing, centrifuge to remove the supernatant and resuspend the pellet in 10 µL protein buffer. Combine equal volumes of the two suspensions and image immediately or after 12 h. Migration efficiency is quantified using Mander’s overlap coefficients (MOC). The coefficients for GFP and BFP are denoted as M_G_ and M_B_, respectively, and are defined as follows:

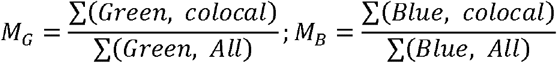

### Comparison of LLPS versus MSE fluorescence intensities

For the liquid-liquid phase separation (LLPS) and MSE groups, GFP was adjusted to a final concentration of 30 μM; the LLPS group was supplemented with 10 % (w/v) PEG 8000, whereas the MSE group received 10 mg/mL silica particles. Both groups were imaged under identical acquisition conditions.

### Radial Intensity Profile Analysis

Radial fluorescence intensity distributions around mineral particles were extracted from raw image stacks. Distances were binned in one-pixel increments (1 µm per bin in this dataset). For each particle, the mean intensity was computed as a function of radial distance r (0–100 µm).

To quantify near-and far-field enrichment, profiles were fitted to a bi-exponential decay model,

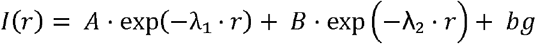

where A,B are amplitude coefficients, λ_1_>λ_2_>0 are the decay constants for the near- and far-field components, and bg is a constant background. Non-linear least squares fitting was performed in MATLAB (lsqcurvefit, R2023a) using bound-constrained optimization. Initial guesses were set to A=0.8·Imax, B=0.2·Imax, λ_1_=20/L, λ_2_=5/L (with L the profile length scale), and bg the minimum observed intensity. Parameters were restricted to non-negative values and the inequality λ _1_>λ _2_ was enforced to ensure physical interpretation.

Goodness of fit was evaluated by the coefficient of determination (R^2^), root-mean-square error (RMSE), and inspection of residual distributions. Approximate 95% confidence intervals were estimated from the Jacobian of the fit, and parameter correlations were reported from the covariance matrix. For all profiles analyzed, the bi-exponential model outperformed single-exponential alternatives (Akaike information criterion), with representative data shown in Fig. 2b (R^2^=0.976).

All plots were generated with custom MATLAB routines, displaying experimental data, overall bi-exponential fit, individual near- and far-field components, and residuals (see Fig. 2b).

To ensure physical interpretability, we performed a post-hoc label reordering on the fitted parameters: the numerically larger decay constant λ was reassigned to λ_1_ (near-field, short-range component) and the smaller to λ_2_ (far-field, long-range component). Because lsqcurvefit does not admit inequality constraints among parameters, we did not enforce λ_1_ > λ_2_ during optimization. Instead, whenever the raw fit returned λ_1_ < λ_2_, we swapped A ↔ B and λ_1_ ↔ λ_2_ in post-processing, aligning the output with the manuscript’s consistent physical labeling. This relabeling leaves the underlying fit unchanged; it merely standardizes the correspondence between numerical values and their physical meaning. **Fluorescence Recovery After Photobleaching (FRAP)** experiments were performed on an Olympus FV4000 laser-scanning confocal microscope using a 100× oil-immersion objective (NA 1.5). GFP-coated SiO_2_ samples had final concentrations of 0.6 mg·mL^-1^ (GFP) and 10 mg·mL^-1^ (SiO_2_). A circular ROI of 2 µm diameter was photobleached with the 488 nm laser at 100% transmissivity for 20 iterations. Recovery images were acquired at 1.4% laser power with 21.2 s intervals for 7.4 min. Fluorescence intensity traces were corrected for background and overall photobleaching, normalized to pre-bleach intensity, and averaged across n = 3 independent particles (mean ± s.d.). Reported recovery values (e.g., ∼20% over 300 s) refer to the plateau of the normalized recovery curve.

### Coupling of Enzyme Cascade Reactions on the Surface of the Same Silica (SiO_2_) Particle

As shown in Figure 3, GOx (final concentration :1 KU/ml, Beyotime Biotechnology), HRP (final concentration:0.2 mg/ml, Beyotime Biotechnology), two kinds of fluorescent proteins (final concentration : of 0.2 mg/ml) were separately added into a centrifuge tube. The mixture was centrifuged at 100 g for 10 seconds. Then, the supernatant was aspirated, and it was washed with 10 μ l of DI water (Solarbio Biotechnology), followed by brief shaking to mix evenly. Then DI water was replaced by 10 μl of substrates solution (glucose with final concentration :1.8 mM, Yuanye Bio-Technology Co., Ltd., and Amplex Red with final concentration : 0.2 μM, Yuanye Bio-Technology Co., Ltd.). Pictures were shot after 20min of mixing.

### Mineral-Surface Templated Compartmentalization through Lipid-Protein Hierarchical Assembly

The lipid used in the experiments related to Figure 4 is palmitoleic acid (Yuanye Bio-Technology Co., Ltd.), and the fluorescent dye for the lipid is Nile red [In Figures 4b-j, the amount of Nile red added is 1 μl (with a concentration of 250 μg/ml) for each case. Yuanye Bio-Technology Co., Ltd.]. In Figures 4a and 4b, 1 μl of palmitoleic acid was added to 10 μl of the silica suspension (with a concentration of 20 mg/ml). In Figures 4d-f, prepare 10 μl of silica particles (with a concentration of 20 mg/ml) coated with GFP (with a concentration of 0.4 mg/ml) according to the procedure in Figure 3b. Then, add 2 μl of palmitoleic acid and gently shake to mix evenly. In Figures 4g-j, prepare lipid-coated silica particles with protein films (GFP or BFP) in the same manner as in Figures 4d-f (using proteins and silica particles of the same concentration, and the amount of lipid used is 1 μl for each case), and then mix the two together. In Figures 4k-m, prepare lipid-coated silica particles with protein films (GFP + GOx or BFP + HRP, with the concentrations of the fluorescent proteins and enzymes being the same as those in Figure 3) respectively in the same manner as in Figures 4d-f. Then, mix the two together and add the substrate solution (with different substrate concentrations the same as those in Figure 3) to prepare a 10 μl suspension. Take photos after 40 minutes of mixing.

### Stochastic simulation of two-dimensional enzyme-cascade kinetics

To evaluate the kinetic consequences of surface confinement, a two-dimensional stochastic simulation was performed (https://github.com/Andyduck-ops/2D-Enzyme-Cascade-MSE). The model tracks individual substrate, intermediate, and product molecules undergoing random diffusion and reaction within a discrete lattice, with catalytic sites corresponding to enzyme I and enzyme II fixed in either bulk or MSE configurations. Reaction probabilities were computed from enzyme-specific catalytic constants and updated at each timestep using Monte Carlo sampling. Simulations were run for over 30,000 time steps with a spatial resolution of 1 nm and a small molecule diffusion coefficient of 1000 nm^2^·s^-1^ (Although significantly smaller than the actual value, it together with the time step dt used in the simulation forms an internally coupled and self-consistent parameter system, here each time step is 0.0015 s, and the total time is 50 s). For the enzyme concentrations at high, medium, and low levels, the values for both bulk and MSE were the same in each case, specifically Enzyme I/Enzyme II = 50/50, 10/10, and 3/3. The resulting product kinetics figure was averaged over 200 replicates to compare catalytic throughput between bulk and surface-confined systems, reproducing the trends shown in Fig. 3c, and the specific parameters are presented in Supplementary Figure 4.

### Introducing RT-PCR into a system containing silica dioxide

Suspend the silica particles with 2X q-RT PCR buffer. Then, following the instructions of the kit (Beyotime Biotechnology), add the different components to the suspension, with the total reaction volume being 20 μL. The final concentration of the template RNA is 14 ng/μL. The sequences of the primers are as follows: 5’-AGAGCTGTTCACTGGTGTCG & 5’-TAGTTGCGTCACCTTCACCC (For CDS of GFP, synthesized by Sangon Biotech).

### Three-dimensional reconstruction of GFP[SiO_2_] and PAL-GFP[SiO_2_] complexes

The prepared samples were imaged with an inverted fluorescence microscope (Nikon Ti2-E, Japan) equipped with a spinning-disk confocal laser-scanner unit (Nova-SD, Airy Technologies Co., Ltd., China) and a 100 × /1.49NA objective (Nikon, Japan). For GFP[SiO_2_] complex, GFP and SiO_2_ at final concentrations of 0.6 mg/mL and 5 mg/mL. For PAL-GFP[SiO_2_] complex, supplement the above mixture with 0.5 µL Rhodamine B dye (10mM in DMSO) and 0.5 µL palmitoleic acid.

### Silica-BFP particles ± Cy5-PAL: permeation assay of Streptavidin-FITC

Prepare 20 µL suspensions of BFP and SiO_2_ at final concentrations of 0.5 mg/mL and 5 mg/mL. After 5 min incubation, centrifuge and discard the supernatant. Resuspend the pellets in 10 µL protein buffer, add 0.1 µL palmitoleic acid containing 10% Cy5-labeled analogue, mix thoroughly, incubate for 5 min, then centrifuge again and remove the supernatant. Finally, resuspend the pellets in 5 µL Streptavidin-FITC solution (Beyotime Biotechnology) and incubate for 30 min.

### Analysis of Fluorescence Intensity

All the raw data were obtained using Fiji (Fiji Is Just ImageJ), and the data visualization was carried out with MATLAB R2023b.

### Figure generation

All figures were created with MedPeer (medpeer.cn).

## Acknowledgements

We express our gratitude to Airy Technologies Co., Ltd., China, for the fluorescence imaging conducted with their spinning-disk confocal microscope (Nova-SD). We express our gratitude to engineer XinYuan Chen of Airy-tech Company for the LSCM operation depicted in Figure 4. We express our gratitude to Hao Xiong, ShiYuan Pei, and Yang Wen for their preparation and reservation of instruments and reagents. We acknowledge Yuzhi Ji and Chen Fan for their assistance during the experiment.

## Supplementary information

## Supplementary Note S1 | Theoretical framework for mineral-guided surface enrichment

### S1.1 Model framework and assumptions

We model the steady-state radial concentration field around a spherical mineral particle of radius *R* immersed in a dilute aqueous solution. The framework couples (i) adsorption– desorption kinetics at the mineral interface (which determine an effective boundary concentration at the particle surface) with (ii) radial diffusion and first-order exchange processes in the diffuse corona. The principal assumptions are:

1. **Hierarchical adsorption**. A dense primary (strongly bound) layer can form at the surface, producing an effective boundary concentration *c*_*surf*_ for a more diffuse secondary corona. Secondary adsorption and intermolecular interactions in the corona create a spatially varying concentration profile extending away from the surface.
2. **Quasi–steady state**. Diffusion and exchange equilibrate on time scales short compared with experimental observation, so that the radial profile may be approximated by a steady solution (time derivatives are neglected).
3. **Linearized corona dynamics**. When concentration deviations in the corona are modest, reaction/exchange terms in the diffuse corona can be approximated by an effective first-order rate *k*_*eff*_. Where nonlinear kinetics are important, the full nonlinear coupling should be solved (see S1.2).
4. **Geometry**. Analyses use the full spherical solution unless explicitly stated; the planar exponential approximation is recovered in the limit *L* ≪ *R* (corona thickness *L* much smaller than particle radius *R*). All symbol units are given in S1.7.

### S1.2 Surface adsorption kinetics and boundary concentration

Primary adsorption at the mineral surface can be described by Langmuir-type kinetics. Let θ(*t*) be the fractional surface coverage; a simple kinetic model reads

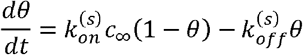

where 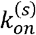 is the adsorption coefficient (units: µM^-1^·s^-1^), 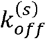 is the desorption rate (s^-1^), and *c*_∞_ is the far-field bulk concentration (µM). At equilibrium,

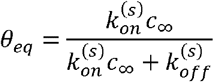

If *c*_*max*_ denotes the maximum surface-associated concentration (µM equivalent for surface-bound material), the effective boundary concentration used in the diffuse-phase calculation is

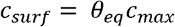

Alternatively, when experimental measurements of surface concentration are available, those values may be used directly in place of the Langmuir estimate.

### S1.3 Steady-state reaction–diffusion and spherical solution

We approximate the diffuse corona by a linearized steady-state reaction–diffusion equation in spherical coordinates. Let ***c***(***r***) denote the concentration at distance ***r*** from the particle center. The linearized steady equation is

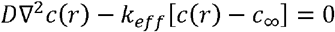

where *D* is the diffusion coefficient (µm^2^·s^-1^) and *k*_*eff*_ is an effective first-order exchange rate (s^-1^). Introducing *λ*^2^ = *k*_*eff*_ /*D* the radially symmetric solution that satisfies *c*(*r* → ∞) = *c*_∞_ and *c* (*R*) = *c*_*surf*_ is

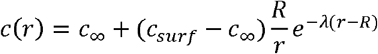

Hence the excess concentration above bulk is

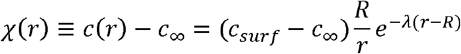

The inverse length *λ* is

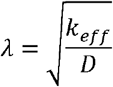

So the penetration length is 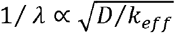.

A practical linearized approximation commonly used for parameter scans is

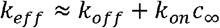

which follows from linearizing adsorption/exchange about the far-field concentration.

When adsorption kinetics or concentrations are strongly nonlinear, the full coupled boundary value problem (Langmuir surface + nonlinear reaction terms) should be solved numerically.

### S1.4 Planar limit and applicability

When the corona thickness *L* satisfies *L* ≪ *R* and the profile is examined for *r* close to *R*, the geometric prefactor *R*/*r* is approximately unity and the spherical solution reduces to the planar exponential:

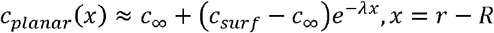

Applying the planar form outside this regime (for example, using the planar expression to examine *r* many times *R* while treating *R* as small) is inconsistent; when *r* ≫ *R* the geometric dilution *R*/*r* must be retained.

### S1.5 Interpretation and recommended parameter scans

Increasing increases penetration length 1/*λ*, thereby extending the radial region where *c*(*r*) may approach or exceed a critical concentration *c*_*crit*_. However, *D* does not directly increase the boundary concentration *c*_*surf*_. Thus whether interfacial LLPS nucleation occurs depends jointly on *c*_*surf*_ and 1/*λ*.

*c*_surf_ can be either set from experimental surface measurements or predicted from Langmuir equilibrium via 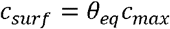. Providing either *c*_*surf*_ or the Langmuir parameters 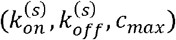 improves reproducibility.

For figures that explore (*r,D*) phase maps (e.g. Fig.2j), we recommend using the spherical solution for all and reserving the planar approximation only when *L* ≪ *R* is explicitly satisfied. To demonstrate robustness, parameter scans should vary *c*_*surf*_, *k*_*on*_, *k*_*off*_ and *D* across experimentally plausible ranges and report the region in parameter space where 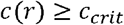.

### S1.6 From concentration field to measured fluorescence — interpretation of exponential fits

In the main text and Supplementary figures (e.g. Fig.2b–c) we represent the measured fluorescence intensity as a sum of two exponential decays plus a background:

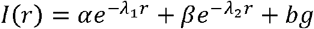

where *I*(*r*) is the measured fluorescence signal at distance from the mineral interface (arbitrary fluorescence units), *α* and *β* are amplitude coefficients, λ_1,2,_ are inverse length scales (µm^-1^), and *bg* is a baseline signal that reflects far-field/bulk contribution and instrument background.

Two interpretational points are important:

#### □Empirical fit vs. mechanistic parameter

The bi-exponential form is an empirical representation that compactly captures a rapidly decaying near-surface component and a more slowly varying tail. A direct mechanistic identification of each exponential with the spherical analytic solution of S1.3 requires caution: the spherical solution predicts a single exponential factor modulated by the geometric prefactor *R/r* under the linearized reaction–diffusion approximation. Therefore, the appearance of two exponentials in the measured intensity may arise from one or more of the following physical scenarios: (i) two distinct exchange regimes (different effective *k*_*eff*_ and/or distinct molecular species with different *D*); (ii) convolution of the concentration profile with the optical point-spread function and instrument response; (iii) coexistence of a primary adsorbed layer plus a diffuse corona with different effective decay; or (iv) nonlinear adsorption coupling not captured by a single linearized *k*_*eff*_.

#### □Mapping λ to model parameters

If independent values for the relevant diffusion coefficient *D* (or for *k*_*eff*_) are available, one may attempt to map a fitted inverse length *λ*_*i*_ to mechanistic parameters using

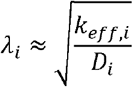

Absent independent *D* (or independent kinetic estimates), the fitted λ_*i*_ should be treated as inverse length scales (units µm^-1^) and reported as characteristic decay lengths 1/*λ*_*I*_ (µm). Any mechanistic mapping must explicitly state the assumed or measured *D* used for that mapping.

**Practical implication for Fig.2b–c:** in all fits we report 1/*λ*_1_ and 1/*λ*_2_ with units µm^-1^ and report 1/*λ*_1,2_ (µm) as characteristic lengths. In the main text we clearly indicate whether these fitted scales are interpreted phenomenologically (length scales only) or are mapped to *k*_*eff*_ using independent/assumed values of (explicitly cited).

### Supplementary Note S2 | Stochastic kinetic simulation of an enzyme-catalyzed cascade reaction under MSE condition

#### Computational Model Framework

We developed a stochastic 2D computational model to simulate enzyme-catalyzed reaction dynamics in both membrane-surface-engineered (MSE) and bulk solution environments. The model implements Brownian diffusion, boundary interactions, and two-step enzymatic reactions (S → I → P) through discrete time-stepping with step size Δt.

#### System Initialization

The simulation begins with particle initialization using the *init_positions* function. For MSE mode, enzymes (GOx and HRP) are uniformly distributed within an annular region defined by particle radius (pr) and film thickness (ft) around a central particle. Substrate particles are randomly placed outside the film boundary region. In bulk mode, all particles are uniformly distributed throughout the simulation box of size L × L. The system maintains separate tracking for substrate, intermediate, and product particles with unique identifiers.

#### Time Evolution Protocol

The simulation progresses through discrete time steps in a loop from t=0 to T_total. Each iteration sequentially executes three core physical processes: Diffusion Step: The diffusion_step function implements heterogeneous Brownian motion. In MSE mode, particles within the film boundary (r ≤ film_boundary_radius) diffuse with coefficient D_film, while those outside use D_bulk. Bulk mode employs uniform D_bulk throughout. All mobile particles (substrate, intermediate, product) undergo independent displacement vectors drawn from a Gaussian distribution with variance 2DΔt.

#### Boundary Handling

The *boundary_reflection* function manages particle interactions with system boundaries. For box boundaries, specular reflection ensures particles remain within the simulation domain. In MSE mode, additional reflection from the central particle prevents penetration by pushing particles to a buffer radius along the normal vector, with special handling for numerical stability near the center.

#### Reaction Kinetics

The *reaction_step* function implements two-stage enzymatic catalysis. Reaction probabilities are calculated using either exact (1-exp(-k_catΔt)) or approximate (k_catΔt) formulations. When enzyme-substrate pairs approach within reaction radii (rGOx + rSub for GOx-S; rHRP + rInt for HRP-I), reactions occur probabilistically with inhibition factors. MSE mode restricts reactions to the membrane ring region. Successful reactions transition particles between states (S→I→P) while setting enzyme busy timers to prevent immediate reuse.

#### Simulation Parameters and Configuration

Key parameters include diffusion coefficients (D_bulk, D_film), catalytic rates (k_cat_GOx, k_cat_HRP), particle radii, film thickness, and simulation box size. The model supports both accurate probabilistic calculations and linear approximations for computational efficiency. Neighbor searching employs KD-tree optimization with GPU acceleration options.

#### Data Collection and Analysis

During simulation execution, the *record_data* function captures particle counts, positions, and reaction events at specified intervals. Output metrics include temporal profiles of product formation, reaction event coordinates, and spatial distributions. The simulation concludes by compiling results into a comprehensive structure containing final particle states and time-course data for subsequent analysis.

This computational framework provides a robust platform for investigating spatiotemporal dynamics of coupled enzymatic reactions in confined and bulk environments, with particular relevance to membrane-based biocatalytic systems.

---

**Supplementary Figure 1.**
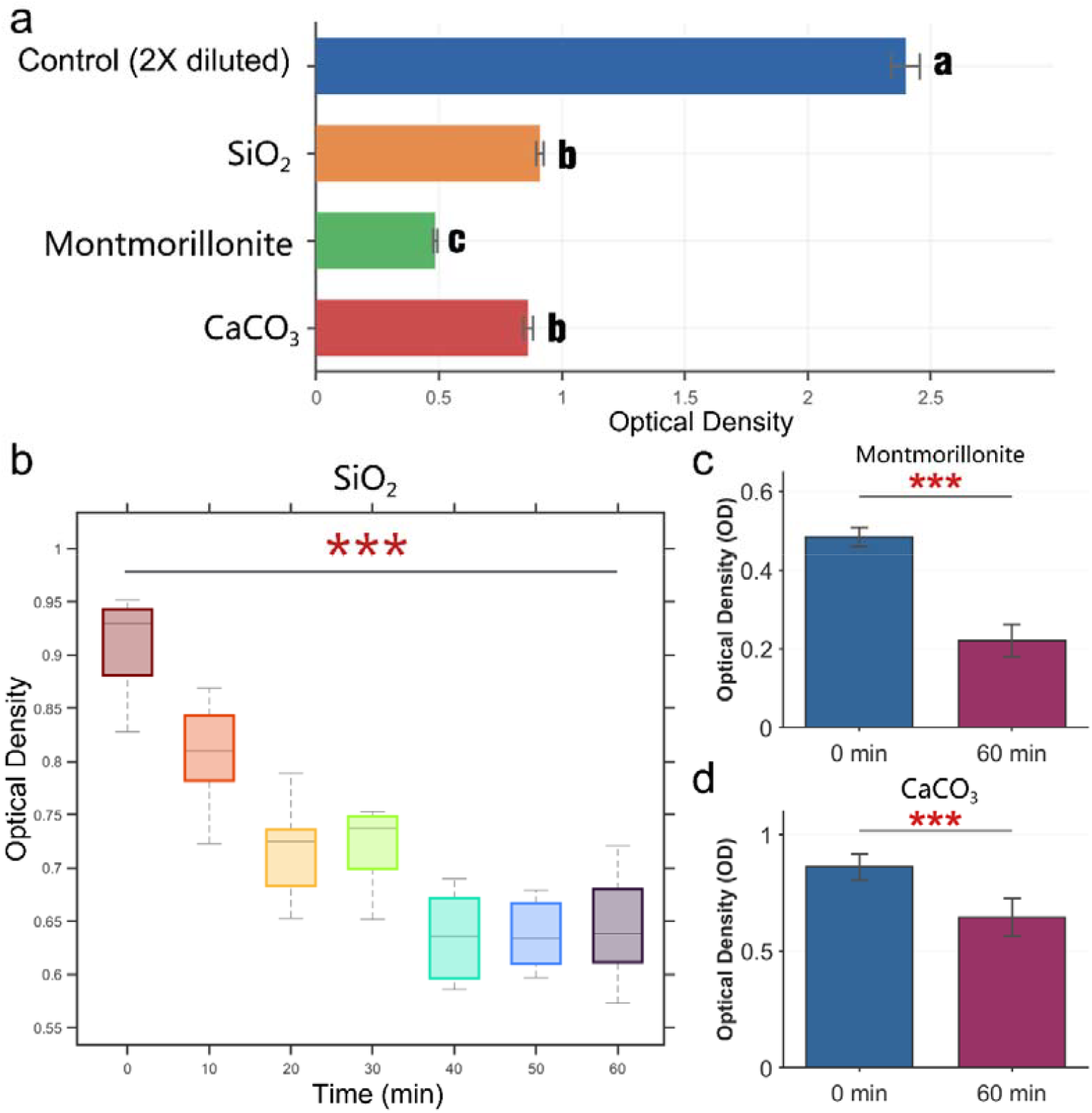
Protein adsorption onto mineral surfaces under elevated temperature. (a) Optical density (OD) of supernatants after 60 min incubation at 70L□°C. Control group: 1 mg·mL^-1^ BSA solution (2 × diluted with buffer); experimental groups: SiO_2_, montmorillonite, or CaCO_3_ mixed with 1 mg·mL^-1^ BSA. (b–d) Temporal OD profiles for SiO_2_ (b), montmorillonite (c), and CaCO_3_ (d) systems. Shorter heating durations (<60 min) were compensated by incubation at ambient temperature. Error bars: s.d. (n = 8); letters and *** denote significant inter-group differences with P < 0.001 (compared to 0-min control). BSA (Beyotime Biotechnology) was dissolved to 1 mg·mL^-1^ in 50 mM Tris-HCl (pH 7.4) buffer. Mineral particles (SiO_2_, montmorillonite, or CaCO_3_) were added to a final concentration of 10 mg·mL^-1^ and incubated with BSA for 60 min at 70□°C unless otherwise noted. Samples were centrifuged to ensure complete precipitation of the minerals (2000g, 30s), and OD of the supernatant was measured at 562 nm using a spectrophotometer (SpectraMax iD3-INJ) and BCA kit (Beyotime Biotechnology).

**Supplementary Figure 3.**
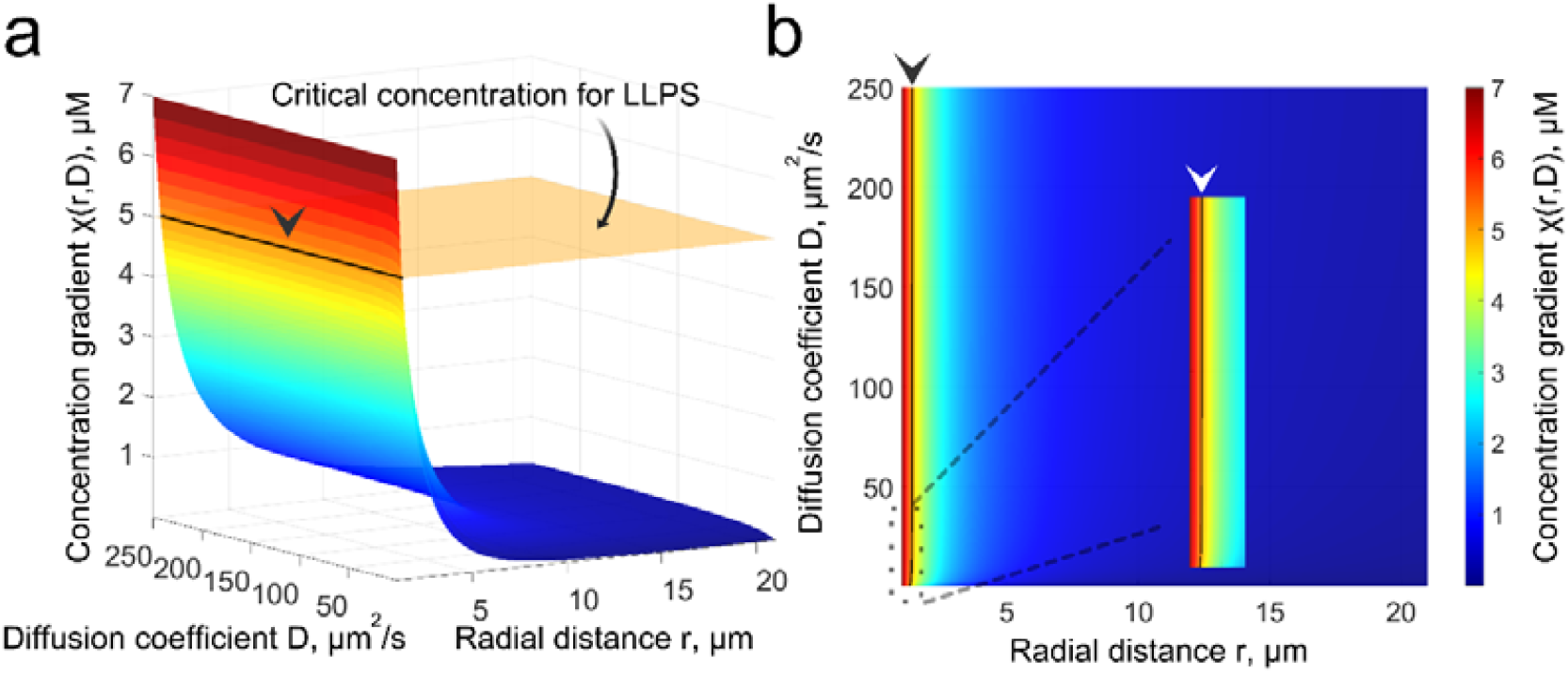
Three-dimensional distribution of the concentration gradient as a function of the diffusion coefficient D and the radial distance r. (a) When, under certain conditions, the local concentration of macromolecules on an inorganic surface exceeds the phase-separation threshold (surface lying above the yellow plane), liquid–liquid phase separation may nucleate at these loci. (b) Top-down view of (a): regions to the left of the solid black line (Indicated by the arrowhead) denote sites where LLPS nucleation is permissible.

**Supplementary Figure 4.**
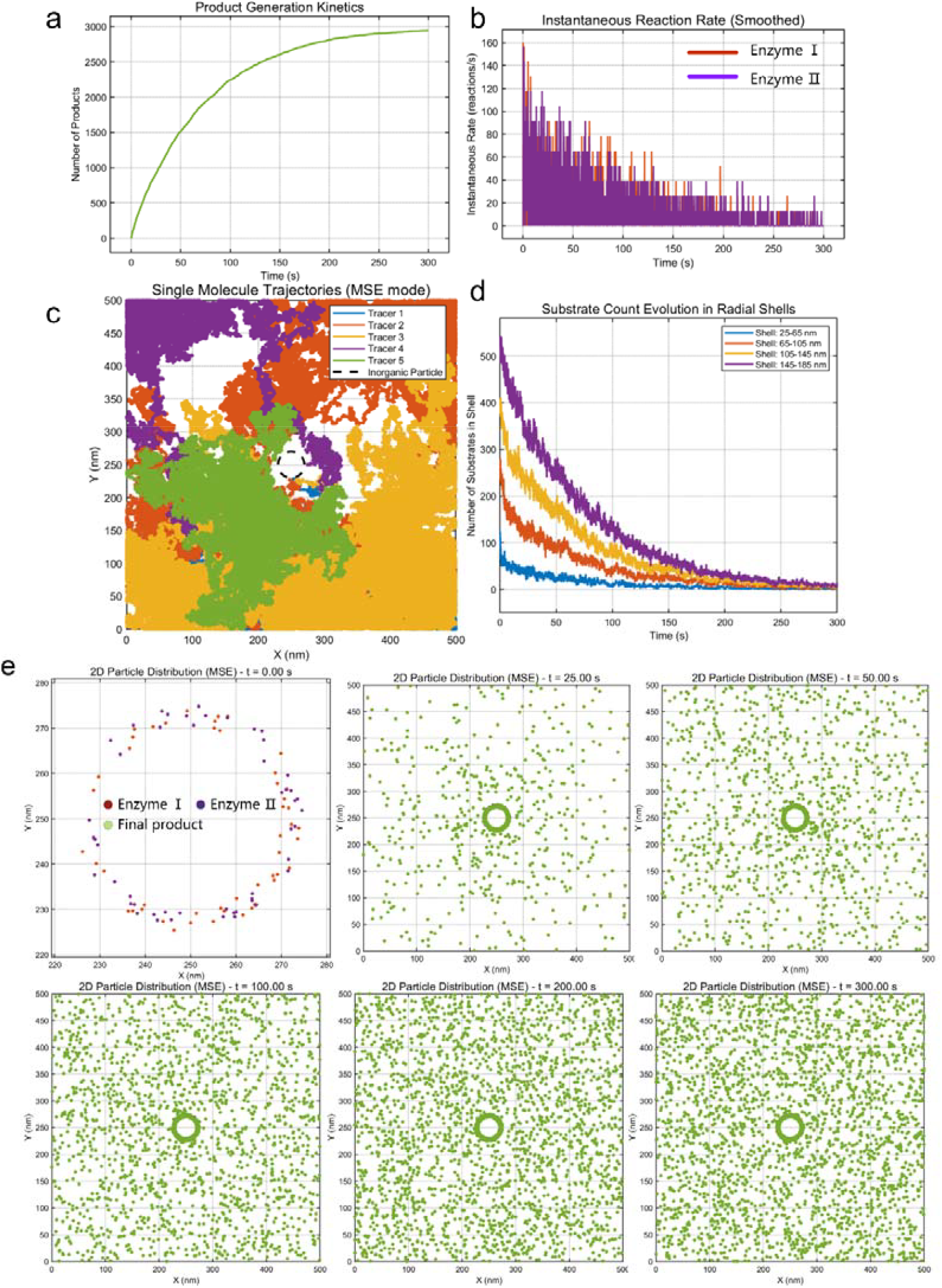
The simulated reaction dynamics of MSE. (a) Cumulative product development curve. The parameters of the simulation code are delineated as follows: The reactive cubic domain possesses an edge length of 500 nm, with a total simulation duration of T = 300 s and a time step of dt = 0.0015 s. The inorganic particle has a diameter of 40 nm. The quantities of the two enzymes are each 50, exhibiting catalytic efficiencies (K_Cat_) of 100 1/s. At t = 0, the substrate count is 3000. The diffusion coefficient for small molecules (substrates, intermediates, and final products) is established at 1000 nm^2^/s; the enzyme film thickness is 5 nm, with enzymes randomly distributed within this film; we assume the diameters of the enzymes are 2 nm, while the diameters of the small molecules are 1 nm. (b) Instantaneous reaction rates of two enzymes, we define it as reactions per second. (c) Independently monitor the motions of 5 small molecules. Any molecule that undergoes conversion—be it from substrate to intermediate or from intermediate to end product—remains classified as a singular small molecule. (d) Quantify the quantity of substrate molecules within uniform shells of 20nm thickness surrounding the surface of the inorganic particle. (e) Snapshots of the reaction progress at successive time points.

**Supplementary Figure 5.**
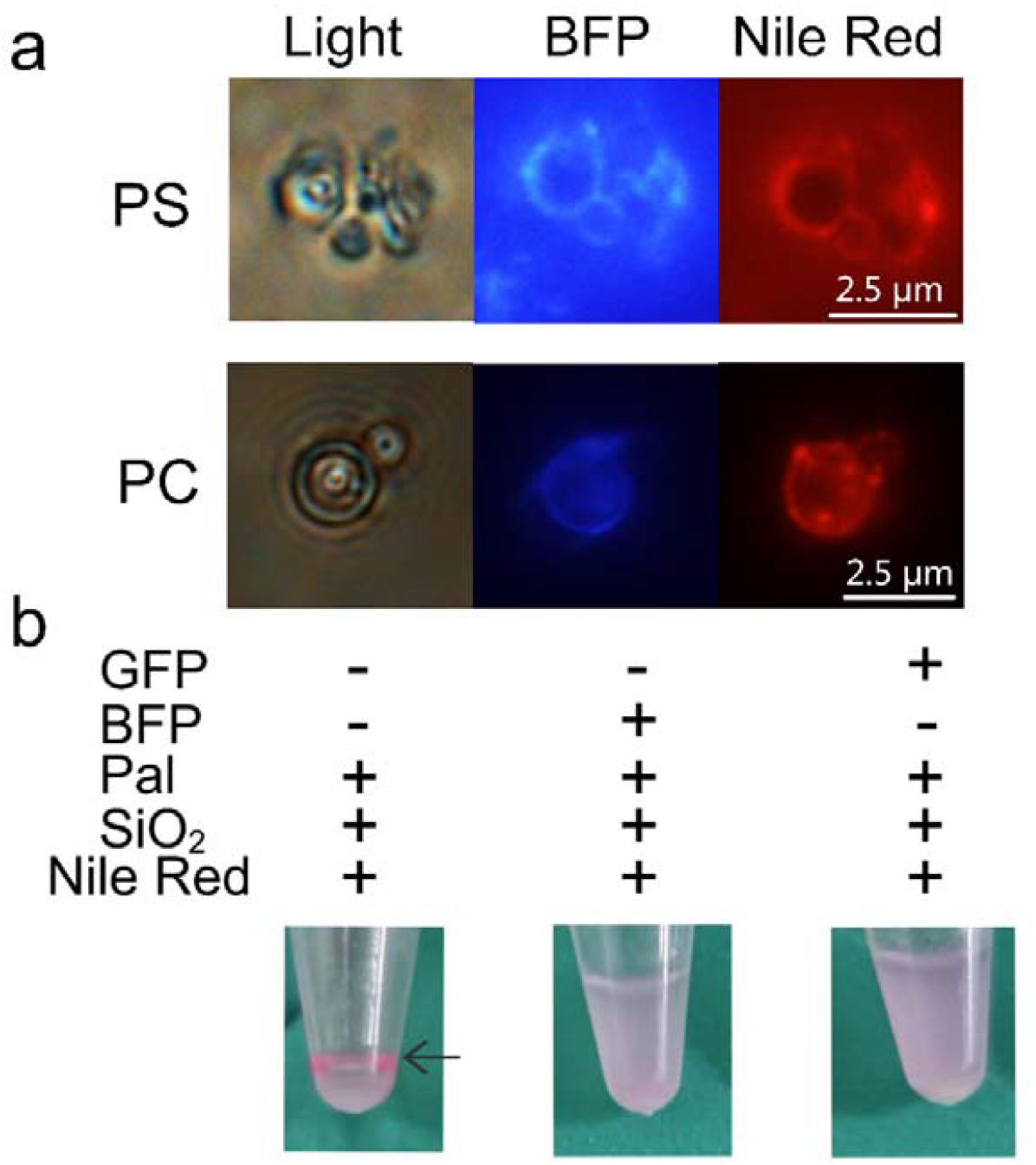
Protein coronas interact with diverse phospholipids and stabilize lipid-mineral suspensions. (a) Co-localization of phosphatidylserine (PS) and phosphatidylcholine (PC) with the protein corona on silica particle surfaces. From left to right: bright-field, BFP-hnRNPA1-IDR, and Nile Red channels. Scale bars: 2.5 µm. (b) This picture illustrates the states of samples comprising palmitoleic acid (Pal), silica (SiO_2_), Nile red (Nile), coupled with the incorporation of BFP and GFP, respectively, following mixing. The left-hand centrifuge tube, containing solely Pal, SiO_2_, and Nile, displays clear stratification (as indicated by the arrow). Conversely, the middle and right-hand centrifuge tubes, upon the incorporation of BFP and GFP respectively, yield suspensions. The three sets of centrifuge tubes depicted in this picture include silica particles at a concentration of 20 mg/ml, palmitoleic acid comprising 1/12 of the total volume, a nile red solution at 250 μg/ml also constituting 1/12 of the total volume, and protein at a concentration of 0.5 mg/ml. Microscopic analysis reveals that the presence of protein modifies the interaction between free lipids and minerals.

**Supplementary table 1.** Parameter definitions and units.

| Parameters | Statements | Units |
| --- | --- | --- |
| $r$ | Radial distance from particle center | $\mu\text{m}$ |
| $R$ | Particle radius | $\mu\text{m}$ |
| $c(r)$ | Concentration distribution function | $\mu\text{M}$ |
| $c_{\infty}$ | Concentration of the bulk phase | $\mu\text{M}$ |
| $c_{surf}$ | Concentration at the mineral surface | $\mu\text{M}$ |
| $c_{max}$ | The maximum surface-associated concentration | $\mu\text{M}$ |
| $D$ | Diffusion coefficient | $\mu\text{m}^2 \cdot \text{s}^{-1}$ |
| $k_{on}$ | Adsorption coefficient used in linearized exchange, for explicit surface adsorption denote $k_{on}^{(s)}$ | $\mu\text{M}^{-1} \cdot \text{s}^{-1}$ |
| $k_{off}$ | Desorption rate | $\text{s}^{-1}$ |
| $k_{eff}$ | Effective first-order rate for the corona, approximated as $k_{off} + k_{on}c_{\infty}$ | $\text{s}^{-1}$ |
| $\lambda$ | $\sqrt{k_{eff}/D}$ | $\mu\text{m}^{-1}$ |

**Supplementary table 2.**
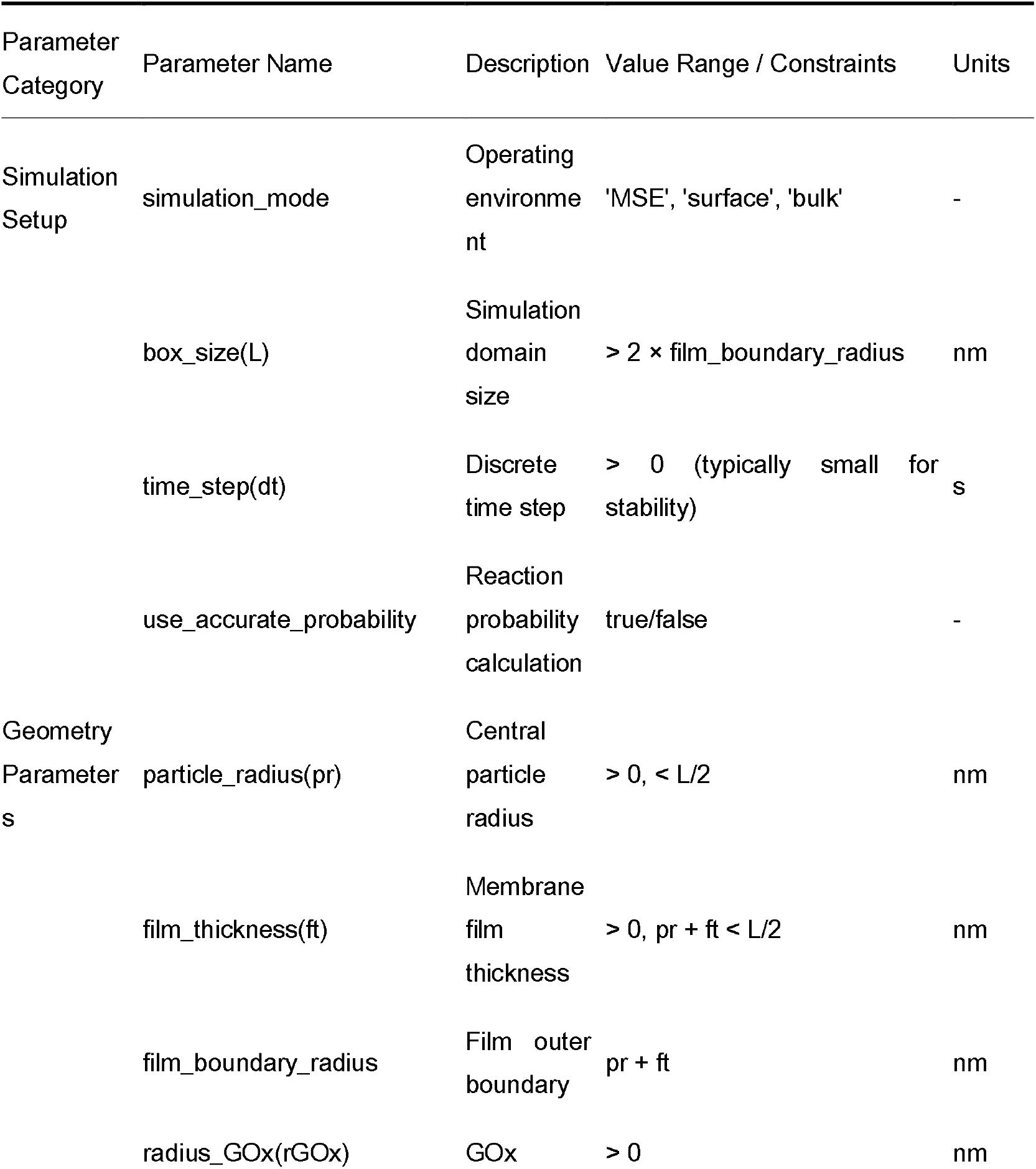
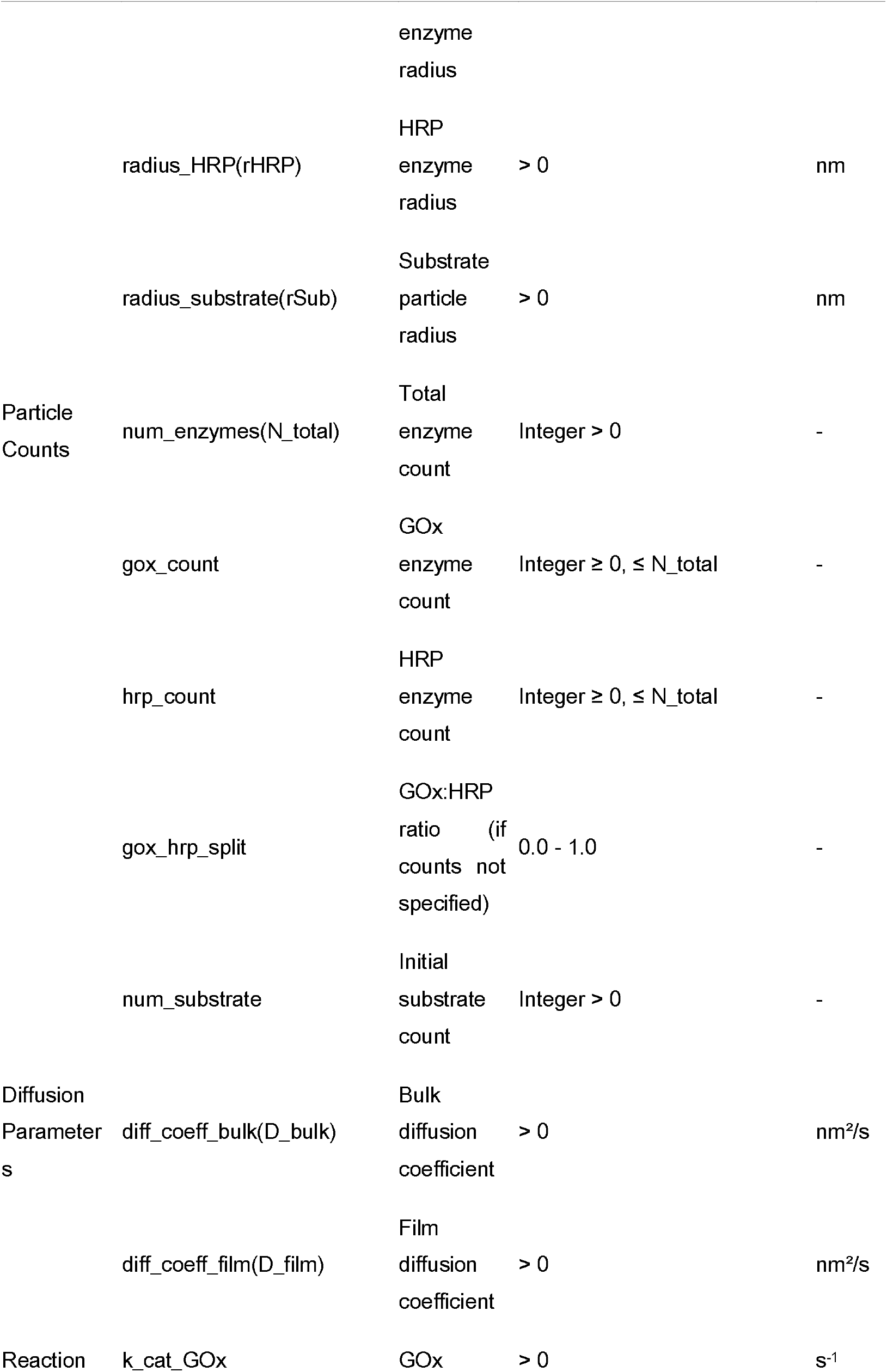
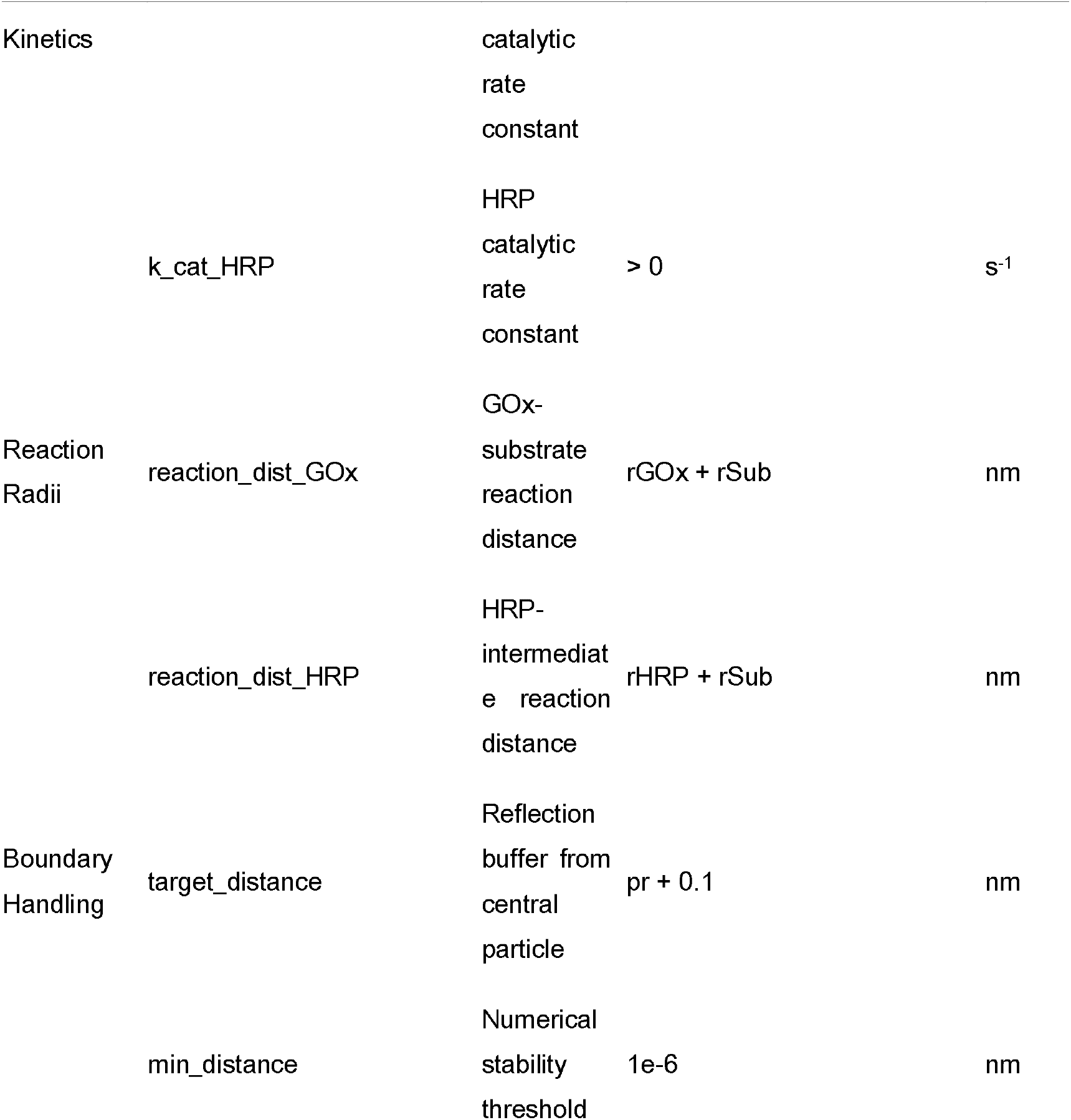
The summary of simulation parameters of an enzyme-catalyzed cascade reaction Summary.

**Supplementary table 3.**
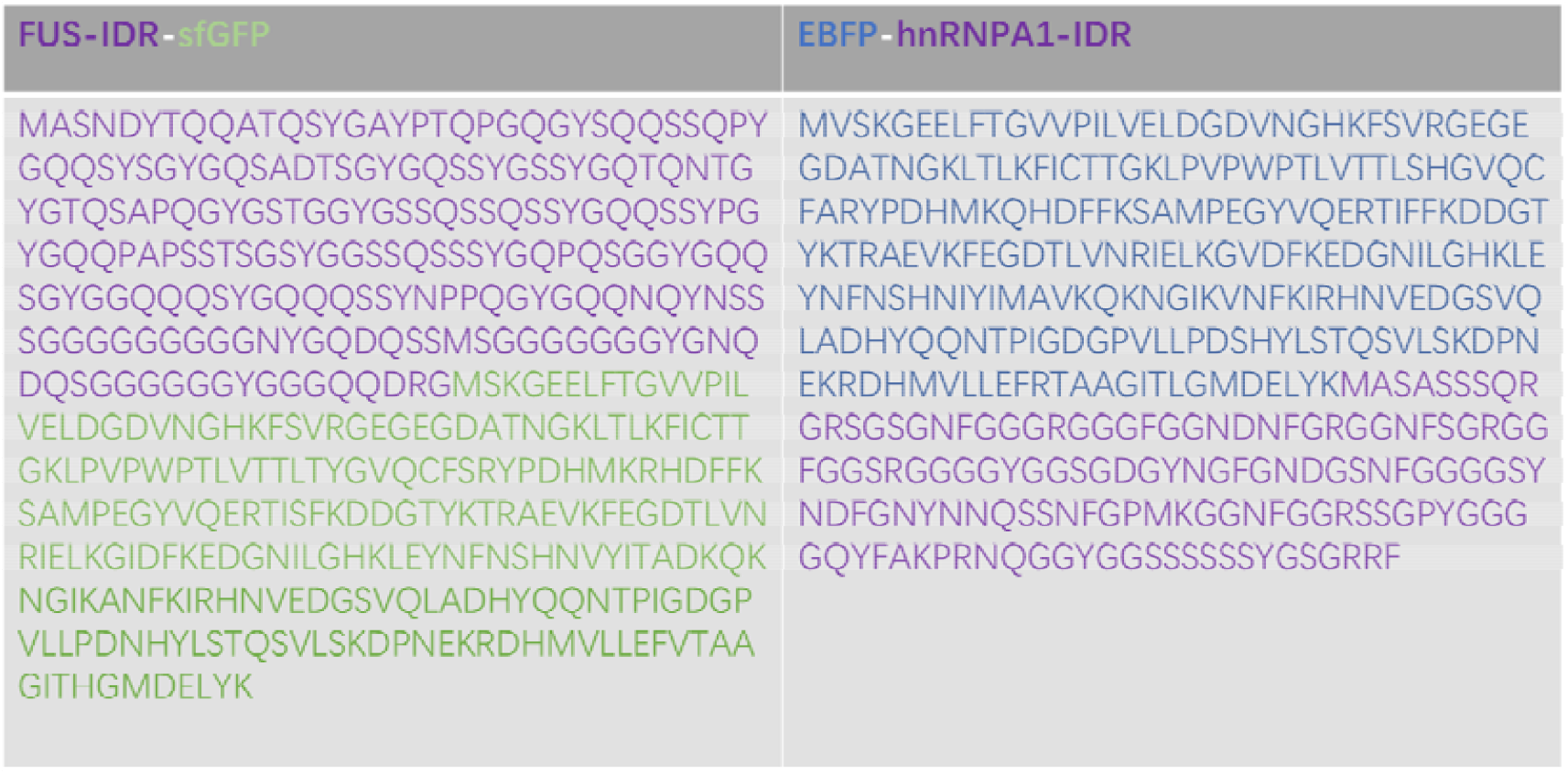
the sequences of two fluorescent proteins The sequences of the two fusion proteins were engineered into the pET-28a(+) plasmid and expressed in the BL21(DE3) strain. This construction strategy was deliberately designed to ensure that the resultant fused proteins harbored a 6×His tag. The cloning sites selected for this manipulation were Ndel and Xhol. The sequence synthesis and plasmid construction were served by Sangon Biotech Company.

**Supplementary table 4.** The composition of non – denaturing lysis/washing/elution/dialysis buffer (pH=8.0).

| Reagents | Final concentration | Source |
| --- | --- | --- |
| NaH<sub>2</sub>PO<sub>4</sub> | 50mM | Beyotime |
| NaCl | 300mM | Beyotime |
| ddH<sub>2</sub>O | n/a | Solarbio |
| NaH<sub>2</sub>PO<sub>4</sub> | 50mM | Beyotime |
| NaCl | 300mM | Beyotime |
| imidazole | 2mM | Beyotime |
| ddH<sub>2</sub>O | n/a | Solarbio |
| NaH<sub>2</sub>PO<sub>4</sub> | 50mM | Beyotime |
| NaCl | 300mM | Beyotime |
| imidazole | 50mM | Beyotime |
| ddH<sub>2</sub>O | n/a | Solarbio |
| 1M Tris-HCl, pH=7.4 | 50mM | Beyotime |
| NaCl | 25mM | Beyotime |
| Glycerol | 10% | YuanYe |
| ddH<sub>2</sub>O | n/a | Solarbio |

